# Sex Differences in Human Subcortical Anatomy are Robust and Replicable in Adulthood, Already Evident Prepubertally, and Aligned with Sex Chromosomal and Gonadal Influences

**DOI:** 10.64898/2026.10.05.756728

**Authors:** Hyo M. Lee, Siyuan Liu, François M. Lalonde, Liv Clasen, Elisa Guma, Alex DeCasien, Dustin Moraczewski, Adam Thomas, Jason P. Lerch, Gabriel A. Devenyi, M. Mallar Chakravarty, Karen F. Berman, Angela Delaney, Armin Raznahan

## Abstract

The existence and mechanistic bases of sex effects on human subcortical anatomy remain unclear. To address these gaps, we combined structural neuroimaging of sex differences in subcortical volume and shape across three independent cohorts (combined N=2,886) with neuroimaging maps of sex chromosomal and endocrine effects from three clinical cohorts (combined N=313, including sex chromosome aneuploidies and isolated gonadotropin-releasing hormone deficiency). We found that human subcortical anatomy shows focal sex differences of moderate-strong effect size during young adulthood (male-biased surface area overlying lateral/ventral putamen and pallidum, nucleus accumbens, ventral posterior complex of thalamus, anterior hippocampus, and medial and central nuclei of amygdala; female-biased surface area overlying caudate head, posterior/dorsal pallidum, mediodorsal, lateral dorsal, lateral geniculate, and medial geniculate nuclei of thalamus, posterior hippocampus, and basolateral and lateral nuclei of amygdala). These regional sex differences show strong within-sample reproducibility at modest sample sizes (N≥200) and are replicable in a fully independent mid-adulthood cohort. Strikingly, we find that the topography of adult subcortical sex differences is already established by pre/early puberty (age 9 years), and the same holds for cortical sex differences. Comparison with neuroimaging data from rare disorder cohorts reveals that subcortical sex differences show a statistically significant spatial and directional congruence with the anatomical effects of X-chromosome dosage in the striatum and effects of testicular hormones in the amygdala. Together, these findings provide a reference map for the fine-grained spatial patterning of sex differences in adult subcortical shape, demonstrating its replicability, establishment before pubertal progression, and spatial concordance with potentially causal sex chromosomal and gonadal effects.

## Introduction

The existence and mechanistic bases of sex differences in human brain organization are topics of intense debate^1,2^. Interest in this question stems from its basic science importance, as well as its potential to shed light on well-established sex differences in diverse brain-related health conditions in psychiatry^3^ and neurology^4^.

Neuroimaging provides our primary means of testing for potential sex differences in the living human brain, with the bulk of data to date stemming from structural neuroimaging^5–12^, and most of these reports focusing on the cerebral cortex^13–16^. Despite mixed results from the initial phase of small and methodologically heterogeneous studies^1,17^, greater access to large datasets and standardized methods has revealed highly reproducible sex differences in regional cortical anatomy above and beyond the male-bias in overall brain size^2,12,18,19^. The potential mechanistic bases of male-female differences encompass sex chromosomal^20^, gonadal^21^, and gender-based experiential factors^22–26^. While biological influences are hard to causally dissect in humans^27^, some sex chromosomal and gonadal effects can be dissociably modelled through naturally occurring medical conditions. By harnessing this strategy, recent work has shown that anatomical sex differences in the cortical sheet are significantly co-localized with anatomical effects of sex chromosome dosage and gonadal function^19^ – helping to prioritize causal models on a region-by-region basis.

The rapid progress in our understanding of neuroimaging sex differences in the human cortex far outstrips that which exists for subcortical structures such as the amygdala, hippocampus, thalamus, pallidum, and striatum. This disparity is particularly striking given the increasingly recognized importance of subcortical structures for most domains of complex human behavior^28–36^. Moreover, subcortical structures are some of the best-established foci of neuroanatomical sex differences in mammalian model systems (especially rodents), with increasingly well-understood mechanistic bases^37–41^. For example, recent neuroimaging studies in mice have expanded on classical histological work^42,43^ to define topographically organized foci of volumetric sex differences across multiple subcortical structures including the amygdala (male-biased in medial, central, and basomedial nuclei), hippocampus (male-biased in dorsal, ventral and proximal CA2, CA3, and dentate gyrus), and thalamus (female-biased in anterolateral, anteromedial, posteromedial, ventral posteromedial, reticular, and mediodorsal nuclei)^39–41^. A large body of rodent experimental research has firmly linked these subcortical sex differences to the action of circulating sex steroids in both perinatal and pubertal brain development^39,41,44,45^ (across all these aforementioned foci), and more recently, to influences of sex chromosome dosage^41^ (e.g., in ventral thalamic nuclei, dorsal hippocampus, and medial amygdala).

Collectively, these studies in rodents – which share the same basic sex chromosomal and gonadal machinery for sex-biased development as humans^46^ – raise the hypothesis that humans too may show localized and reproducible sex differences in subcortical anatomy that can manifest before puberty, respect the topography of subcortical connectivity, and overlap with anatomical influences of sex chromosome dosage and gonadal function. However, several gaps in knowledge and longstanding technical obstacles in human research have left these major hypotheses largely untested.

First, studies of sex-biased subcortical anatomy in humans have tended to focus on the bulk volume of each subcortical structure^2,12,13^ rather than regional effects within each structure. The handful of studies on regional effects have yielded varying results^7,20,47^ and have not directly quantified the effect size of any sex differences observed, nor their reproducibility and replicability. Second, while there is some evidence from longitudinal neuroimaging studies at lower MRI field strengths that sex differences in regional subcortical anatomy are highly dynamic over puberty^20,47^, it remains unclear if puberty is critical for establishing the regional distribution of sex differences in the adult human brain, or whether it acts to reinforce earlier-established effects (as has been seen for the mouse amygdala^39^). This question carries major consequences for the developmental timing of mechanistic models for sex-biased brain anatomy. Finally, we have so far lacked studies that can test the potential role of sex chromosomal and gonadal factors in shaping subcortical anatomical sex differences in humans, despite emerging methods for addressing this question using rare disorders of sex chromosome dosage (sex chromosome aneuploidies [SCAs]) and endocrine functioning (isolated gonadotropin-releasing hormone deficiency [IGD])^19^.

To address these aforementioned knowledge gaps and technical hurdles, we applied computational modeling of subcortical shape^48,49^ to structural MRI data from three large cohorts of typical controls (combined N=2,886) and clinical cohorts of SCAs and IGD (combined N=313). Our goals were to: (i) fine-map adult sex differences in striatum, pallidum, thalamus, hippocampus, and amygdala; (ii) characterize their reproducibility as a function of sample sizes and replicability in an independent adult cohort; (iii) determine the extent to which they are already evident in pre/early pubertal childhood; and (iv) establish whether they are spatially aligned with the congruent anatomical effects of sex chromosome dosage and gonadal hormone production.

## Results

### Sex Differences in Adult Subcortical Shape Follow a Multi-Focal Pattern that is Obscured by Analysis of Bulk Volume

We analyzed structural MRI scans from the Human Connectome Project Young Adult^50^ (HCP-YA) S1200 release (N = 1,085, aged 22-37 years, 592 females; **Sup Table 1**) to characterize sex differences in subcortical bulk volume and regional shape in early adulthood. All scans were processed through the MAGeT-Brain pipeline^48,49^ (**Figure 1A, left**) to segment five subcortical structures (striatum, pallidum, thalamus, hippocampus, and amygdala) for measurement of bulk volume (**Figure 1A, middle**) and regional shape (**Figure 1A, right**). Following quality control, we applied linear models to estimate the standardized effect size and statistical significance of sex differences while controlling for effects of age and global size on the volume and shape of each subcortical structure (**Methods**).

**Figure 1.**
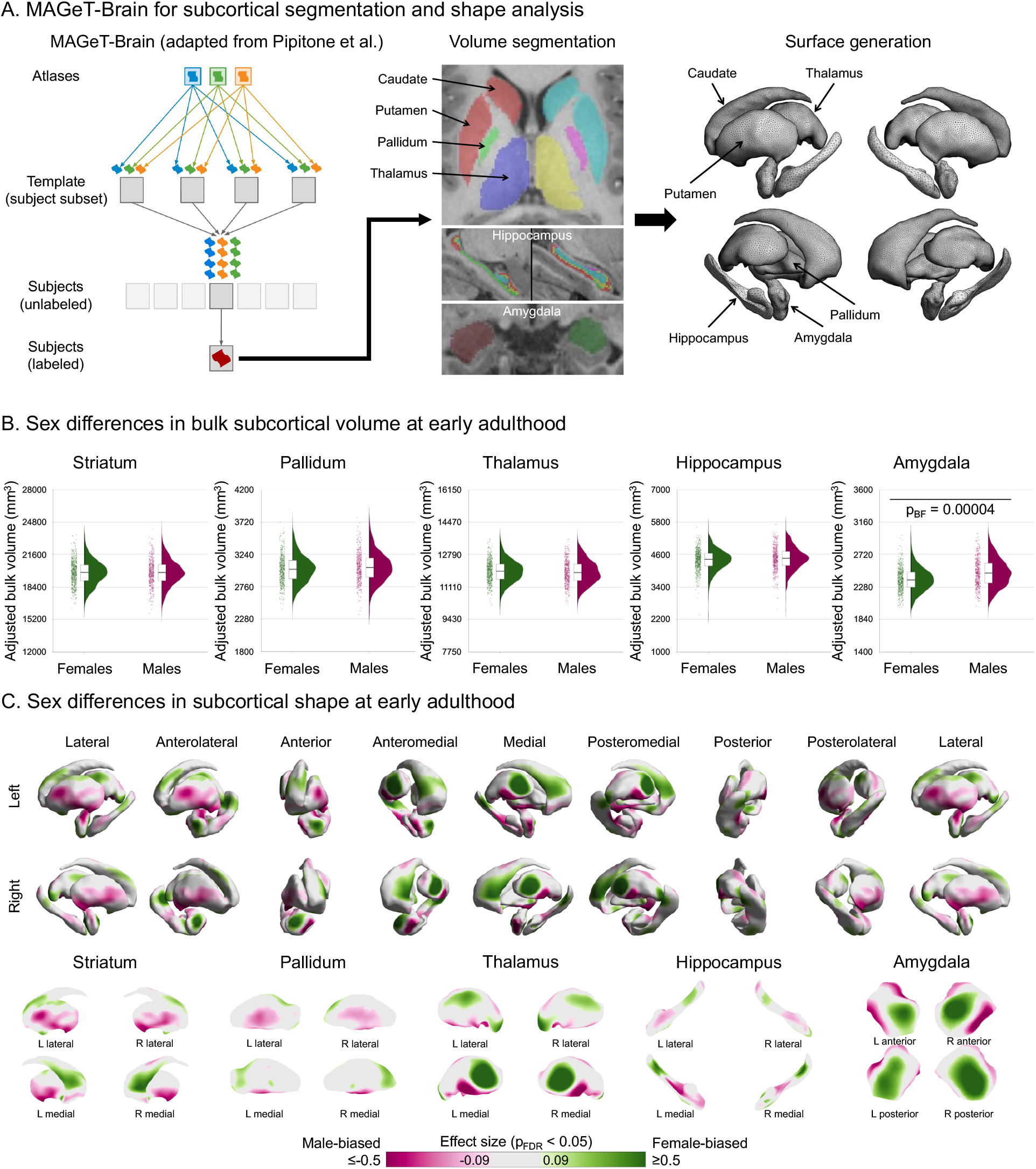
Resolving sex differences in subcortical volume and shape at early adulthood. **A.** MAGeT-Brain transforms a small subset of high-quality, manually segmented atlases to a representative subset of subjects to serve as study-specific templates, which are propagated to each subject to generate multiple candidate segmentations that undergo majority voting to determine the final label (left, adapted from Pipitone et al., 2014). This procedure yields volumetric segmentation for striatum, pallidum, thalamus, hippocampus, and amygdala (middle). The reference surfaces for these structures in the atlases are warped to subject space to calculate vertex-wise surface area (right). Arrows pointing to the five subcortical structures indicate their names. **B.** Bulk volume of subcortical structures (adjusted for age and total brain volume) for females (green) and males (magenta). The boxes, histograms, and jittered dots indicate medians bounded by first and third quartiles, distribution, and individual values. Horizontal bars denote structures showing significant sex differences at Bonferroni-corrected t-test p_BF_ < 0.05. **C.** Standardized effect sizes for sex differences in regional surface area (controlling for age and total surface area) showing statistically significant female-biases (green) and male-biases (magenta) based on a two-tailed t-test followed by FDR correction at p_FDR_ < 0.05 across vertices on both hemispheres within each structure. The top row shows the five subcortical structures together in rotating views. The bottom row shows each structure separately. M: males, F: females, L: left, R: right. Source data are provided in Source Data - Sex Differences in Human Subcortical Anatomy.xlsx.

Statistically significant sex differences in bulk volume after correction for total brain volume were found only in the amygdala (standardized effect size |β| = 0.33, p_BF_ = 0.00004 after Bonferroni correction for multiple comparisons across 5 subcortical structures; **Figure 1B, Sup Figure 1A**). The remaining structures showed small and statistically non-significant sex differences in volume (|β| ≤ 0.11, p_BF_ > 0.05). However, all 5 subcortical structures showed statistically significant sex differences in regional shape when controlling for their overall size (**Figure 1C** surviving false discovery rate [FDR] correction for multiple comparisons across vertices; **Sup Figure 2A** unthresholded effect size maps).

Sex differences in subcortical shape were spatially extensive and included moderate to large effect sizes: 50.6% of the striatal surface was statistically significantly sex-biased in shape with |β|: 0.09-0.54; pallidum (37.8%, |β|: 0.10-0.49); thalamus (56.9%, |β|: 0.09-0.82); hippocampus (55.4%, |β|: 0.11-0.61); and amygdala (71.4%, |β|: 0.10-0.65). Prominent foci of significantly sex-biased surface area included the ventral putamen and distal dorsal caudate tail, lateral and ventral pallidum, ventral thalamus, anterior hippocampus, and dorsolateral, medial, and ventromedial amygdala (all male-biased), as well as the dorsal putamen, caudate (head, body, and tail), posterior and dorsal pallidum, dorsomedial, dorsolateral, and posterior-ventral thalamus, posterior hippocampus, and anterior and posterior amygdala (all female-biased). We verified the robustness of these sex differences in subcortical shape to alternative (allometric) methods for control of overall structure size, and also confirmed the expected pattern of globally male-biased regional surface area in the absence of control for structure size (**Sup Figure 3**).

These results provide a fine-grained reference map for the complex topography of sex differences in human subcortical anatomy during young adulthood – underlining the value of shape-based vs. purely volumetric analysis^2,13^.

### Adult Sex Differences in Subcortical Shape Show Strong Within-Sample Reproducibility in a Sample Size Dependent Manner

To assess the within-sample reproducibility of sex differences in subcortical shape, we quantified the spatial similarity of sex differences estimated between two non-overlapping split-halves of the HCP-YA sample, repeated 10,000 times and at varying split-half sizes (**Figure 2A**). For example, in each repetition and at the split-half size of 450, sex differences were recomputed in a split-half sample of 450 (225 females and 225 males) and the other split-half sample of 450 (225 females and 225 males), and the Pearson correlation between the two resulting unthresholded effect size maps was estimated. For all five structures, we observed that the median spatial correlation of sex differences increased with increasing split-half sample size, asymptotically reaching a plateau at split-half size of ∼200. At the maximum split-half sample size of 450 for the HCP-YA dataset, we found strong reproducibility indicated by the median and interquartile range (IQR) of spatial correlations for each subcortical structure: striatum (0.84 [0.80-0.87]), pallidum (0.78 [0.73-0.82]), thalamus (0.89 [0.85-0.91]), hippocampus (0.85 [0.81-0.88]), and amygdala (0.92 [0.89-0.93]). These findings: (i) show that sex differences in regional shape are a reliable morphological property of the human brain; (ii) inform sample size considerations for future studies; and (iii) help to interpret observed spatial instabilities of reported sex differences across historically smaller sized studies^1^.

**Figure 2.**
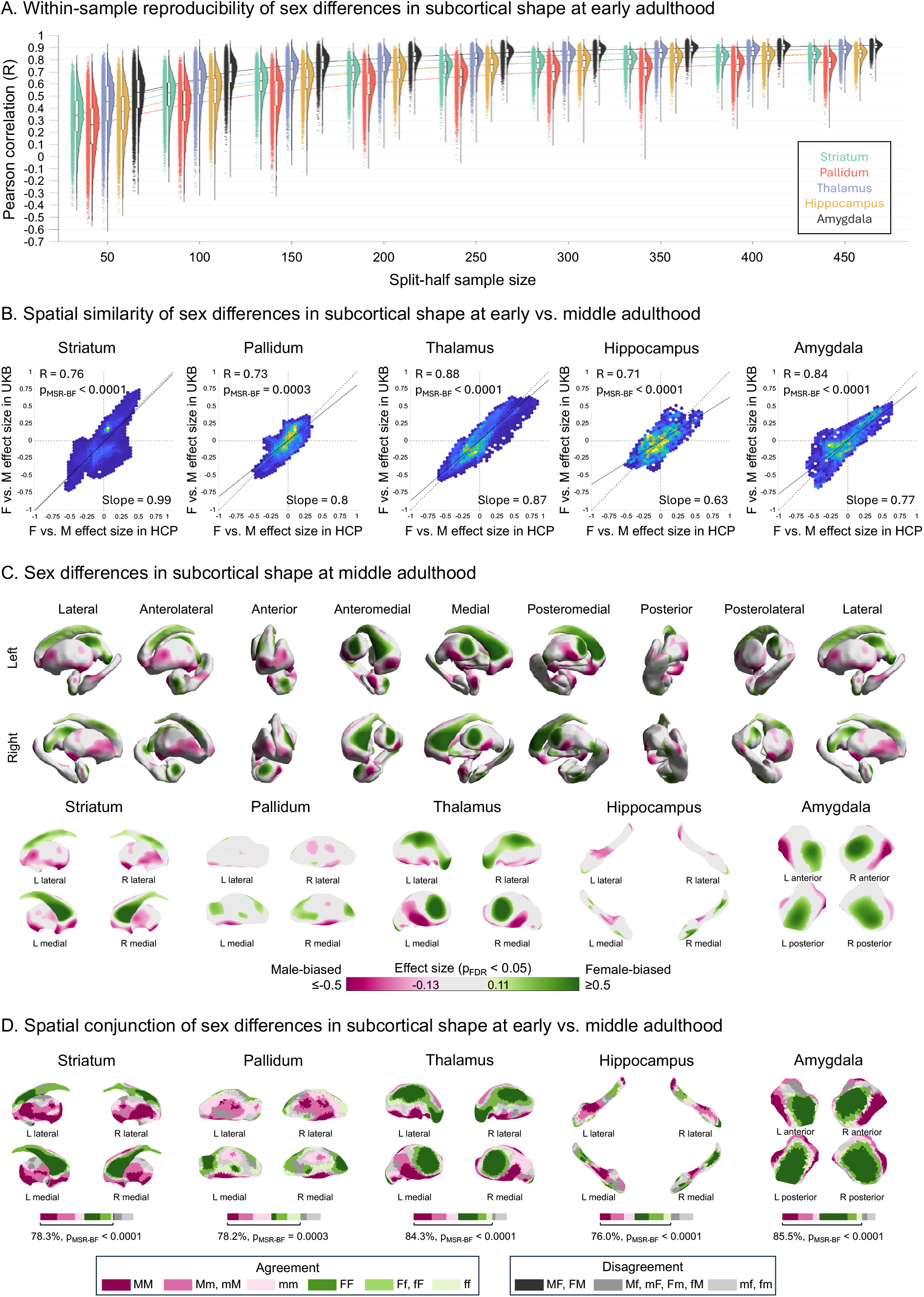
Reproducibility and replicability of sex differences in adult human subcortical shape. **A.** Structure-wide spatial correlations (R) of sex differences in regional surface area of striatum (green), pallidum (red), thalamus (purple), hippocampus (yellow), and amygdala (black) across 10,000 instances of sex-balanced split-halves of varying sizes (50 to 450 in increments of 50) of young adults within the HCP-YA dataset. In each instance, sex differences were recomputed in each independent split-half and correlated between halves. The boxes, histograms, and jittered dots indicate medians bounded by first and third quartiles, distribution, and individual values for R. **B.** Scatter plots showing structure-wide spatial correlations for the sex differences in regional surface area at early vs. middle adulthood (i.e., between HCP-YA and UKB-MA datasets). The vertical and horizontal dotted lines are x = 0 and y = 0 lines. The diagonal dotted line indicates y = x identity between the two datasets. The solid line indicates the regression line predicting mid-adulthood sex differences from those in young adulthood. Legends show Pearson correlation (R), Bonferroni-corrected one-tailed statistical significances against a spatial null model based on 10,000 Moran spectral randomizations of HCP-YA sex differences (**Methods**), and slope of the regression line. **C.** Standardized effect sizes for sex differences in regional surface area (controlling for age, total surface area, and site) showing statistically significant female-biases (green) and male-biases (magenta) based on a two-tailed t-test followed by FDR correction at p_FDR_ < 0.05 across vertices on both hemispheres within each structure at middle adulthood (UKB-MA). The top row shows the five subcortical structures together in rotating views. The bottom row shows each structure separately. **D**. Conjunction maps showing 3 levels of agreement between statistically significant anatomical sex differences in the HCP-YA and UKB-MA datasets. Green and magenta regions are concordantly female- and male-biased between datasets, respectively (dark: both datasets significant in the same direction; intermediate: either significant in the same direction; light: both non-significant in the same direction). Gray regions are discordantly sex-biased between datasets (dark: statistically significant in opposite directions between datasets; intermediate: opposite directions between datasets and statistically significant in one; light: opposite directions between datasets, but statistically non-significant in either). Bars show the areal proportion of subcortical surface falling within each of these conjunction categories. Brackets and percentages indicate observed areal proportion of agreement between the two datasets. p_MSR-BF_ indicates the statistical significance of concordant areal proportion using a one-tailed spatial permutation test based on 10,000 Moran spectral randomizations of HCP-YA sex differences (**Methods**). M: males, F: females, L: left, R: right. Source data are provided in Source Data - Sex Differences in Human Subcortical Anatomy.xlsx.

### Sex Differences in Subcortical Shape at Early Adulthood are Highly Replicable in Middle Adulthood

We quantified the out-of-sample replicability of sex differences in subcortical shape in early adulthood against those computed using identical image processing pipeline and statistical models controlling for age, total brain size, and site (**Methods**) in an independent cohort from mid-adulthood collected using different MRI scanners in a different country (UK Biobank^51^, n = 669, 375 females, 5-year age band of 45-50 years closest to HCP-YA representing mid adulthood, henceforth “UKB-MA”; **Sup Table 1**).

At the bulk level, consistent with analysis in early adulthood (**Figure 1B**, **Sup Figure 1A**), we found male-biased amygdala after correction for sex differences in overall brain size (standardized effect size |β| = 0.31, p_BF_ = 0.00001 after Bonferroni correction for multiple comparisons across 5 subcortical structures; **Sup Figure 1B**). However, the mid-adulthood cohort also showed significant female-biases in striatal and pallidal volume (|β| = 0.24 and 0.21, p_BF_ = 0.002 and 0.024, respectively), which had not been apparent in early adulthood, while the thalamus and hippocampus remained non-sex-biased (|β| ≤ 0.06, p > 0.05).

At the regional level, we computed the standardized effect size maps for sex differences in subcortical shape in middle adulthood (**Sup Figure 2B**). These regional sex differences followed a very similar spatial pattern to those seen in early adulthood—as quantified by structure-wide Pearson R correlations for sex effects of 0.76 (striatum), 0.73 (pallidum), 0.88 (thalamus), 0.71 (hippocampus), and 0.84 (amygdala). These correlations all survived testing for Bonferroni-corrected statistical significance against a spatial null model based on 10,000 Moran spectral randomizations^52,53^ (p_MSR-BF_ < 0.0005; **Figure 2B**, **Methods**). The regression slopes for these spatial relationships were furthest below 1 for the hippocampus (0.63) and amygdala (0.77), indicating a strongly preserved topography, with an attenuated magnitude of sex differences in the shape of these structures in mid vs. early adulthood.

We next localized those sex differences in mid-adulthood that survive FDR correction across vertices on both hemispheres within each subcortical structure at p_FDR_ < 0.05 (**Figure 2C**). The areal proportion of significance and magnitude of these sex differences were similar to those in early adulthood (**Figure 1C**): 60.5% of the striatal surface was sex-biased in shape in mid-adulthood with |β|: 0.11-0.72; pallidum (30.3%, |β|: 0.15-0.53); thalamus (52.0%, |β|: 0.13-0.63); hippocampus (37.0%, |β|: 0.14-0.50); and amygdala (53.4%, |β|: 0.15-0.67). Replicability of direction and statistical significance of sex differences between early and middle adulthood was formalized by conjunction analyses (**Methods**), which showed agreements for ≥76% of the surface of all 5 structures (**Figure 2D**; p_MSR-BF_ ≤ 0.0003 for all structures; **Methods**).

Thus, sex differences in human subcortical shape are highly reproducible within, and replicable between, adult samples – identifying several foci of statistically significant sex biases in subcortical surface area that: include moderate to large effect sizes; are detectable and reproducible in early adulthood; and are still apparent in an independent mid-adulthood cohort 20 years older in mean age.

### The Topography of Adult Sex Differences in Subcortical Shape is Already Apparent at Pre/Early Puberty, and this Same Phenomenon is Seen for Cortical Anatomy

To test if the spatial distribution of sex differences in adult subcortical shape (as assayed in the HCP cohort) is dependent on puberty, we applied identical image processing pipeline and statistical models used for analyses of adult subcortical anatomy (**Methods**) to an independent cohort of pre/early pubertal 9-year-olds from the Adolescent Brain Cognitive Development (ABCD-9Y) Release 2.0.0 (N = 1,132, aged 9-9.5 years, 569 females; **Sup Table 1**).

As context for these regional analyses, we first considered the bulk volume of each subcortical structure (controlling for overall brain size), and found that sex differences in pre/early puberty were restricted to male-biases in the amygdala (standardized effect size |β| = 0.13, p_BF_ = 0.021; **Sup Figure 1C**), presaging the larger male-bias observed in adulthood. However, moving beyond analyses of bulk volume (**Methods**) revealed – for all 5 subcortical structures examined – that the spatial patterning of unthresholded regional anatomical sex differences in the pre/early pubertal ABCD sample was highly similar to the independent HCP adult sample (**Sup Figure 2A vs. 2C,** Pearson R correlations of 0.7 [striatum], 0.59 [pallidum], 0.51 [thalamus], 0.68 [hippocampus], and 0.66 [amygdala]; all p_MSR-BF_ < 0.05; **Figure 3A**; **Methods**). The regression slope for prediction of regional sex differences in adult subcortical shape from those in pre/early-puberty was notably greater than 1 for the amygdala (1.82), suggesting that this structure shows an especially prominent pubertal accentuation of sex differences in its shape (regression slopes for other structures: 1.01 [striatum], 0.66 [pallidum], 0.93 [thalamus], and 1.2 [hippocampus]).

**Figure 3.**
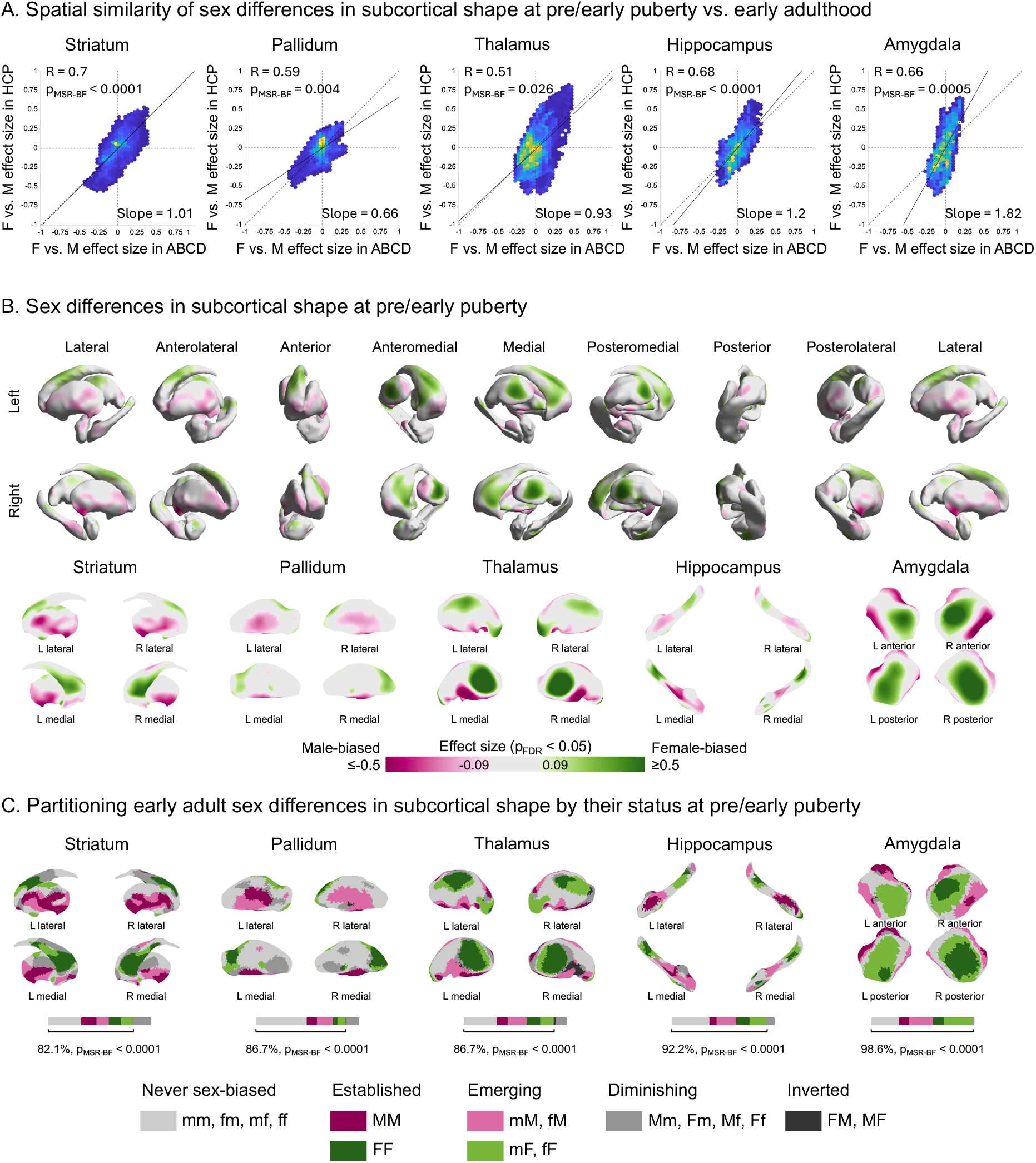
The topography of adult sex differences in subcortical shape is already apparent in pre/early puberty. **A.** Scatter plots showing structure-wide spatial correlations for the sex differences in regional surface area between pre/early puberty (ABCD-9Y) and early adulthood (HCP-YA). The vertical and horizontal dotted lines are x = 0 and y = 0 lines. The diagonal dotted line indicates y = x identity between the two datasets. The solid line indicates the regression line. Legends show Pearson correlation (R), Bonferroni-corrected one-tailed statistical significances against a spatial null model based on 10,000 Moran spectral randomizations of HCP-YA sex differences (**Methods**), and slope of the regression line predicting HCP-YA sex differences based on ABCD-9Y sex differences. **B.** Standardized effect sizes for sex differences in regional surface area (controlling for age, total surface area, and site) showing statistically significant female-biases (green) and male-biases (magenta) based on a two-tailed t-test followed by FDR correction at p_FDR_ < 0.05 across vertices on both hemispheres within each structure in pre/early puberty (ABCD-9Y). The top row shows the five subcortical structures together in rotating views. The bottom row shows each structure separately. **C**. Conjunction maps showing the partitioning of anatomical sex differences in early adulthood (HCP-YA) by their status in pre/early puberty (ABCD-9Y). “Never sex-biased” is defined as regions where sex-biases are non-significant at both pre/early puberty and early adulthood (lightest gray). “Established” is defined as regions where female- and male-biases are significant in the same direction in pre/early puberty and early adulthood (darker green and magenta). “Emerging” is defined as regions where female- and male-biases are significant in early adulthood but non-significant in pre/early puberty (lighter green and magenta). “Diminishing” is defined as regions where female- and male-biases are significant in pre/early puberty but non-significant in early adulthood (medium gray). “Inverted” is defined as regions where female- and male-biases are significantly in the opposite directions between pre/early puberty and early adulthood (darkest gray). Bars show the areal proportion of subcortical surface falling within each of these conjunction categories. Brackets and percentages indicate areal proportion of consistent (never sex-biased, established, and emerging) sex differences between pre/early puberty and early adulthood. p_MSR-BF_ indicates the statistical significance of consistent areal proportion using a one-tailed spatial permutation test based on 10,000 Moran spectral randomizations of HCP-YA sex differences (**Methods**). M: males, F: females, L: left, R: right. Source data are provided in Source Data - Sex Differences in Human Subcortical Anatomy.xlsx.

Correction for multiple comparisons across vertices (p_FDR_ < 0.05) revealed that several foci of statistically significant sex differences in subcortical shape are already apparent in pre/early puberty (**Figure 3B**), with prominent examples in the caudate head and mediodorsal thalamus (female-biased) as well as the ventral putamen and thalamus and overlying the central and medial nuclei of the amygdala (male-biased). These foci showed a slightly smaller spatial extent and magnitude than those in adulthood (**Figures 1C**, **2C**) (pre/early pubertal spatial extent and absolute effect size ranges: striatum [44.8%, |β|: 0.09-0.44]; pallidum [27.4%, |β|: 0.10-0.43]; thalamus [36.7%, |β|: 0.10-0.48]; hippocampus [27.7%, |β|: 0.11-0.31]; and amygdala [21.6%, |β|: 0.11-0.24]). A conjunction analysis of statistically significant sex differences (**Methods**, **Figure 3C**) revealed that foci of sex-biased subcortical shape that are already established in pre/early puberty typically lie within or adjacent to the larger sex-biased terrains that are apparent in adulthood, with most remaining regions of sex-biased subcortical shape in early adulthood already showing the same directional bias in pre/early puberty (**Figure 3C).** Amongst those regions of sex-biased subcortical anatomy that are established by the start of puberty and persist into adulthood, the greatest disparities in effect size (larger in adulthood) were seen in the dorsomedial, ventromedial, and posteroventral thalamus and the basal, medial, and dorsal amygdala (**Sup Figure 5A**). We also observed large swaths of each subcortical structure that were consistent for their lack of a statistically significant sex difference in shape in both pre/early puberty and early adulthood (**Figure 3**, light gray regions). Taken together, these patterns of consistency – already showing a statistically significant or directionally congruent sex difference in pre/early puberty to statistically significant sex differences in adulthood, or not showing a statistically significant sex difference at either age point – accounted for the vast majority of all subcortical surface vertices (striatum [82.1%], pallidum [86.7%], thalamus [86.7%], hippocampus [92.2%], and amygdala [98.6%]). These areal proportions were significantly elevated against a spatial null model based on 10,000 Moran spectral randomizations of early adult sex differences followed by Bonferroni correction across the five structures (p_MSR-BF_ < 0.0001; **Methods**).

Given the striking observation that core foci of sex-biased subcortical anatomy in human adulthood appear to already be in place by pre/early puberty, we tested and confirmed that the same phenomenon holds for sex differences in cortical anatomy (**Methods, Sup Figures 4 and 5B**). Foci of statistically significant sex differences in adult cortical anatomy that are already apparent and statistically significant at pre/early puberty include: primary somatosensory cortex, auditory cortices, dorsolateral, lateral, and medial prefrontal cortices, dorsal parietal cortex, and parahippocampal cortices (all female-biased) as well as insular cortex, opercular area, angular gyrus, fusiform complex, orbitofrontal and ventrolateral prefrontal cortices, retrosplenial complex, hippocampus, and entorhinal cortex (all male-biased).

Taken together, these results reveal that the basic regional topography of anatomical sex differences in adulthood – for both subcortical and cortical anatomy – can already be recovered in unrelated cohorts of pre/early pubertal 9-year-olds. This observation strongly suggests that the genetic and/or environmental factors driving sex differences in adult brain anatomy must (at least in part) be operative before puberty.

### Foci of Sex-Biased Subcortical Shape are Conspicuously Sensitive to Variations in X-Chromosome Dosage and Testicular Hormone Production

To test potential candidate mechanistic hypotheses for sex differences in human subcortical anatomy, we asked if sex-biased subcortical regions tend to overlap with subcortical regions that are also sensitive to effects of sex chromosome dosage and hypothalamic-pituitary-gonadal axis function, which are the two dominant biological sources for sex-biased development in humans and all other placental mammals^44,45^.

We modelled sex chromosome dosage effects on subcortical shape by comparing neuroimaging data in XXY and XYY male patients against matched euploidic XY controls (Total N = 272; 99 XXY males and 92 XY controls; 34 XYY males and 47 XY controls, **Sup Table 1**). These conditions occur in 1/576 and 1/851 live male births^54^, respectively, and prior work at low MRI field strengths has suggested they induce substantial alterations in subcortical shape above and beyond their effects on total brain size^20,55^. While XXY syndrome is associated with secondary endocrine abnormalities and treatments that could potentially alter subcortical anatomy, supernumerary X-chromosome dosage effects have been shown to be consistent amongst multiple SCAs irrespective of their gonadal status (i.e., XXY, XXX, XXXXY)^55^ and mirrored in X-monosomy^56,57^. These reports suggest that their anatomical effects are primarily reflective of direct sex chromosome dosage influences on the brain.

Similarly, we modelled gonadal influences on human subcortical anatomy by harnessing a neuroimaging dataset in males with IGD (estimated prevalence 1 in 8,000 to 30,000 boys^58,59^) and healthy male controls (N=41, 19 IGD, **Sup Table 1**). In brief, IGD is characterized by hypogonadism due to a congenital deficit in hypothalamic drive for pituitary release of the gonadotropins that are required to drive typical testicular development and function^60^. Consequently, compared to controls, males with IGD tend to experience lower levels of circulating testicular steroid hormones during perinatal periods shown to be essential for testosterone-dependent masculinization of the mammalian brain^59^ (although the severity of perinatal hypogonadism can vary across individuals^61^). This period of hypogonadism is extended for IGD patients who are not diagnosed and treated with steroid replacement until later childhood and adolescence^62^. Given this heterogeneity in postnatal steroid treatment, the contrast between XY control males vs. males with IGD primarily captures perinatal influences of intact testicular hormone production on subcortical anatomy.

We used identical imaging and analytic procedures to those employed for analysis of normative sex differences in subcortical anatomy (**Methods**) to estimate standardized effect sizes for X-chromosome dosage, Y-chromosome dosage, and testicular hormone production on the volume and shape of each subcortical structure. We then characterized the congruence of these three effects with the normative sex differences observed at young adulthood.

For brain-size adjusted measures of bulk subcortical volume, we found that increasing X- and Y-chromosome dosage both induced significant volume reductions (**Sup Figure 6A and 6B**) for the pallidum (X: p_BF_ = 0.00003; Y: p_BF_ = 0.0007), hippocampus (X: p_BF_ = 0.03; Y: nominal p = 0.049), and amygdala (X: p_BF_ = 0.000002; Y: p_BF_ = 0.0007), with additional associations between increasing X-chromosome dosage and reduced striatum volume (p_BF_ = 0.0005). Conversely, testicular hormone production effects were non-significant for all five structures (**Sup Figure 6C**). Considering the sex chromosome complements in typical females and males, X-chromosome dosage contributions to volume reductions of amygdala, striatum, and pallidum are congruent with the observed normative male-biases in these three structures (**Sup Figure 1A, 1B and 1C**).

We then generated standardized effect size maps for X-chromosome dosage (i.e., XXY vs. XY, termed “X”), Y-chromosome dosage (XYY vs. XY, termed “Y”), and testicular hormone production (XY controls vs. IGD XY, termed “T”; **Sup Figures 7A, 7B, and 7C**) on regional subcortical shape, and identified those effects that survived FDR correction across vertices on both hemispheres within each subcortical structure at p_FDR_ < 0.05 for X and Y or nominal thresholding at p < 0.05 for T (**Figures 4A, 4B, 4C, top row**). The X and Y effects maps replicated several features of those described in smaller independent cohorts of XXY and XYY individuals scanned with lower field-strength MRI^20,55^, specifically for caudate tail, nucleus accumbens, putamen, pallidum, dorsolateral and dorsomedial thalamus, and posterior hippocampus. The T map, which to our knowledge is the only available description of early hypogonadism effects on human subcortical anatomy, revealed a nominal association of intact early testosterone production with ventromedial caudate, caudate body, ventral pallidum, dorsolateral thalamus, anterior hippocampus, and regions overlying central and medial nuclei of the amygdala.

**Figure 4.**
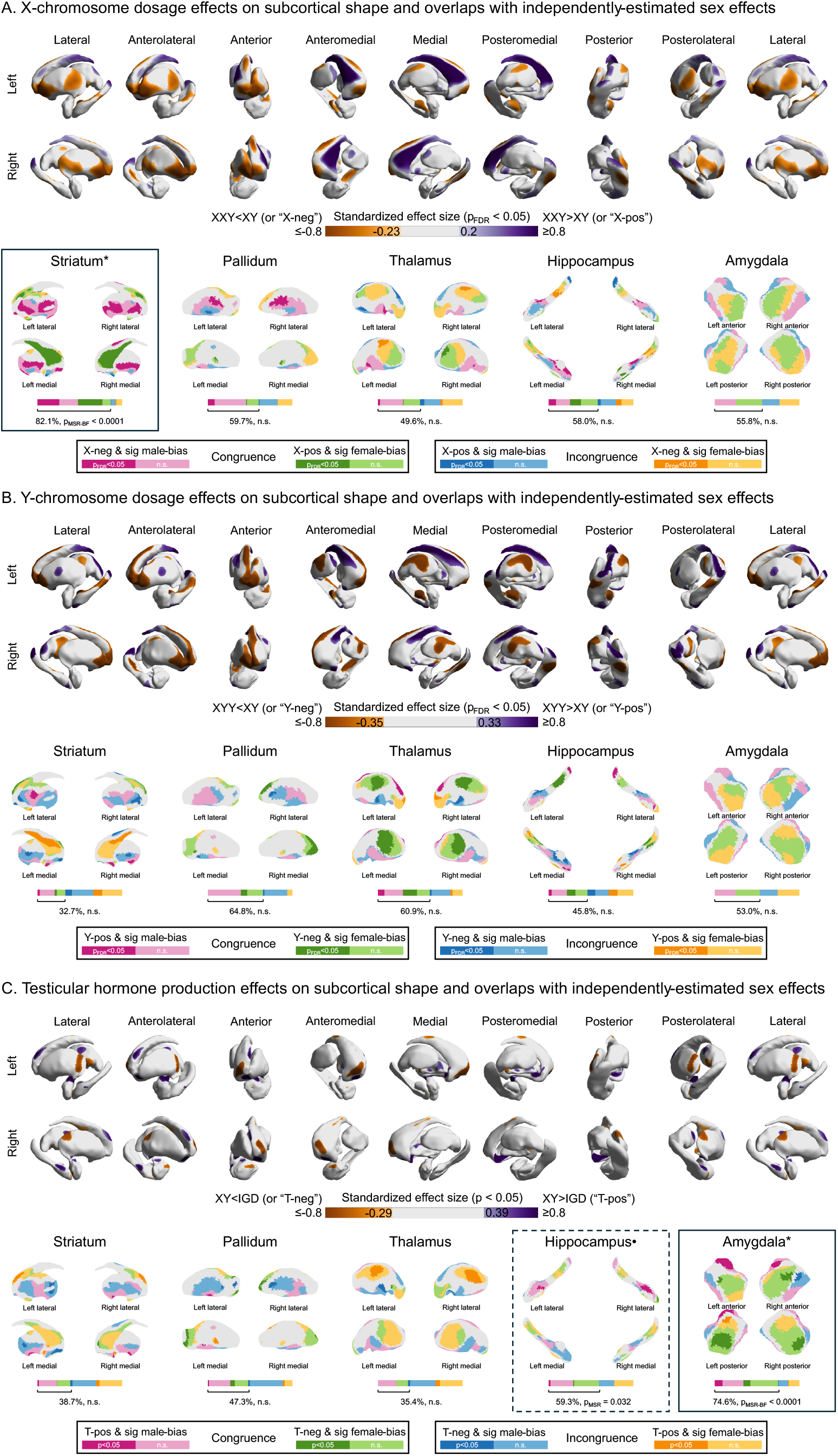
Effects of sex chromosome dosage and testicular hormone production on regional subcortical shape and overlaps of these effects with maps of sex-biased subcortical shape in early adulthood. **A.** Standardized effect sizes for X-chromosome dosage effects (XXY vs. XY) on regional surface area (controlling for age and total surface area) that are statistically significant based on a two-tailed t-test followed by FDR correction at p_FDR_ < 0.05 across vertices on both hemispheres within each structure. Positive effect sizes (XXY>XY or “X-pos”) are shown in purple, while negative effect sizes (XXY<XY or “X-neg”) are shown in orange. Bottom row shows vertex-wise congruence of adult sex differences from the HCP-YA dataset with the effects of X-chromosome dosage. **B.** Same as panel A but for Y-chromosome dosage effects (XYY vs. XY). **C.** Same as panel A but for effects of intact testicular hormone production (male controls vs. males with IGD). For all panels, congruence is defined when X-pos, Y-neg, or T-neg overlaps with significant female-bias (green) and when X-neg, Y-pos, or T-pos overlaps with significant male-bias (magenta). Incongruence is defined when X-neg, Y-pos, or T-pos overlaps with significant female-bias (orange) and when X-pos, Y-neg, or T-neg overlaps with significant male-bias (blue). Darker colors indicate when X, Y, or T effect is significant, while lighter colors indicate when it is non-significant. The bars show the areal proportions at which these conjunction cases make up the subcortical surface. Brackets and percentages indicate the congruent areal proportions. p_MSR-BF_ indicates statistical significance of congruent areal proportion using a one-tailed spatial permutation test based on Moran spectral randomizations of HCP-YA sex differences (**Methods**). Boxes and * indicate statistical significance at p_MSR-BF_ < 0.05. Dashed box and • indicate nominal significance at p_MSR_ < 0.05. Source data are provided in Source Data - Sex Differences in Human Subcortical Anatomy.xlsx.

By generating conjunction maps (**Figures 4A, 4B, 4C, bottom row**) of X, Y, and T effects (**Figures 4A, 4B, 4C, top row**) on subcortical shape with those of normative sex effects (**Figure 1C**), we could classify sex-biased subcortical regions according to the directional (in)congruence of co-localized X, Y and T effects (**Methods**). This conjunctional algorithm was identical to that applied to the cortical sheet in prior work^19^. We define congruent effects as those where the direction of X, Y, and T effects is compatible with the direction of normative sex differences (i.e., where female-biased: XXY>XY, XYY<XY, or control<IGD; where male-biased: XXY<XY, XYY>XY, or control>IGD). We then differentiated between “hard” congruences (where X, Y, and T effects reached statistical significance; **Figures 4A, 4B, 4C, top row**) and “soft” congruences (where X, Y, and T effects fell below statistical significance, **Sup Figures 7A, 7B, 7C**). These analyses revealed two instances of statistically significant spatial congruence (observed congruent overlaps vs. a spatial null model based on 10,000 Moran spectral randomizations of early adult sex differences followed by Bonferroni correction across the five structures [p_MSR-BF_ < 0.0001; **Methods**]): (i) X-dosage and sex effects on the striatum, and (ii) T effects and sex effects on the amygdala (solid black boxes, **Figures 4A, 4C**). Significant X-dosage and sex effects were directionally congruent for 82.1% of the striatal surface (including ventral putamen, distal dorsal caudate tail, caudate head and tail). Nominally significant T effects and significant sex effects were directionally congruent for 74.6% of the amygdalar surface (regions overlying central and medial nuclei of the amygdala). Post-hoc analyses indicated that X-dosage effects on striatal surface area and T effects on amygdalar surface area were strongest for foci of significant female- and male-biased area (respectively) that are already apparent in pre/early puberty (**Sup Figure 8**). These findings suggest that these potential causal influences were more influential on early established vs. puberty-dependent sex differences within each structure.

## Discussion

Our study advances understanding of sex-biased human subcortical organization along three key directions.

First, we establish that humans show a highly reproducible and replicable spatial pattern of sex differences in adult subcortical anatomy. These localized effects – detected through surface-based analysis of regional subcortical shape – exist in all five structures studied herein, regardless of whether the structure shows a sex difference in brain-size adjusted volume. Localized sex differences possess a rich spatial organization that coheres with known topographies of connectivity within the subcortex and beyond^63^. For example, we see matching sex effects at neighboring facets of different structures despite each structure having been analyzed separately (e.g., male-biased area at ventromedial putamen, ventral thalamus, anterior hippocampus, and medial amygdala; female-biased area at dorsolateral thalamus and medial caudate head and body). Regional subcortical sex differences also often echo those previously described for the corresponding cortical regions^19^ to which they are most densely connected (e.g., male-biased medial amygdala, anterior cingulate, anterior insula, and piriform cortex form canonical circuits underlying social, survival, and reproductive behaviors^64,65^; female-biased dorsomedial thalamus, precuneus, and anterior medial prefrontal cortex form key components of the default mode network^66^). Strikingly, several of the localized sex effects we detect in humans show matching sex effects at equivalent subcortical regions in mice, including those overlying lateral geniculate, medial geniculate, and mediodorsal nuclei of thalamus and caudate head (female-biased in both species) as well as those overlying medial and central nuclei of the amygdala and anterior hippocampus (male-biased in both species)^40,41^. Given that mice lack the complex interweaving of sex and gender effects seen in humans, these conserved sex differences may index sex chromosomal and/or endocrine aspects of sex that are shared between species. However, it is important to avoid deterministic accounts for the human sex differences we describe herein. These sex differences all involve large distributional overlaps between males and females; are as of yet unclear in their functional significance^67^; and are consistent with causal contributions from any mix of sex chromosomal, endocrine, gender-based, or other environmental influences^22–25,68,69^. Nevertheless, our findings clearly establish a robust and replicable topography of sex differences in human subcortical anatomy, which coheres with known topographies of brain connectivity and replicates several subcortical sex differences described in mice.

Second, we show that the reproducible topography of sex differences in adult subcortical anatomy is already apparent in pre/early-puberty, and that this same phenomenon of early establishment is also seen for regional sex differences in cortical anatomy. These observations stem from comparison of adult sex differences with those in an independent cohort of 9-year-olds who are verified as being prepubertal or only at the very earliest stages of puberty by both physical maturation and pubertal hormone levels^70^. Comparing the magnitude of effect sizes between cohorts reveals that adult sex differences are an accentuation of regional effects already established before pubertal progression. Thus, it appears that the basic topography of neuroanatomical sex differences in adulthood must be established at some point between organogenesis and late childhood – with subsequent pubertal amplification (especially in mediodorsal, ventral posterior complex, and lateral/medial geniculate nuclei of thalamus, basolateral, lateral, central, and medial nuclei of amygdala, insula, and subgenual/anterior cingulate cortex). This observation helps to narrow the temporal window for operation of whatever genetic and/or environmental factors shape the topography of subcortical sex differences.

Third, we establish that, for the striatum and amygdala, regional sex differences in subcortical shape are significantly aligned (both spatially and directionally) with independently estimated effects of X-chromosome dosage (as estimated through comparison of XXY and XY individuals) and intact testicular function (as estimated through comparison of controls and individuals with IGD). These findings in the subcortex extend our earlier observation of the same alignment of sex effects with X-dosage and testicular effects on regional cortical anatomy^19^. Distinguishing congruent vs. incongruent overlaps of sex effects with those of sex chromosome dosage and testicular function helps to narrow mechanistic hypotheses for regional sex differences. However, colocalization is not proof of causation, and there are assumptions in the use of sex chromosome aneuploidies and IGD to proxy mechanisms of interest. Some of these assumptions are empirically supported: animal models indicate that sex chromosome dosage effects on anatomy in aneuploidy align with those in euploidy^41^; although X-chromosome aneuploidy effects on brain may theoretically reflect secondary endocrine effects in males, comparison with other sex chromosome aneuploidy effects suggests they primarily index chromosome dosage effects^71^; for males with IGD, despite variable neonatal phenotype (micropenis and cryptorchidism vs. normal), pubertal history (absent vs. partially spontaneous), and variable treatment, the vast majority, if not all, share a degree of perinatal hypogonadism^72,73^. Nevertheless, future work would benefit from access to larger neuroimaging samples with rare SCAs and endocrine disorders (allowing better testing for source of variance) and triangulation with complementary study designs for estimating sex chromosomal (e.g., statistical genetics^74^) and gonadal (e.g., puberty^75^, menstrual cycle^76–78^, pregnancy^79,80^, menopause^81,82^, and andropause^83,84^) effects within typically developing cohorts.

Notwithstanding these cautions in mechanistic interpretation, our findings establish a statistically surprising convergence between sex effects on subcortical anatomy and those of putatively causal X-chromosome dosage and testicular function effects. Our observations for the amygdala are especially notable for highlighting regions overlying the medial amygdala as showing a prepubertally established anatomical male-bias that becomes larger by early adulthood and overlaps with areas that become smaller in males with IGD. This constellation of findings aligns closely with developmental and experimental studies of sex-biased amygdala anatomy in mice, where an early postnatal volumetric male-bias is established through masculinizing effects of testosterone^85–87^, with a later accentuation^39,88^ of this male bias through the pubertal testosterone surge. Together, these observations strongly suggest that testosterone plays a role in establishing early male-biases in size of the medial nucleus of the amygdala in humans^89^, which are then elaborated by the pubertal surge in testosterone^21^.

Taken together, our study establishes that adult sex-biases in human subcortical shape have a multi-focal topography that is highly stable across independent adult cohorts, already evident in pre/early puberty (a phenomenon which we establish also holds for sex differences in cortical anatomy), and congruently aligned with X-chromosomal and gonadal influences. For the amygdala in particular, resulting mechanistic hypotheses in humans closely match proven mechanistic drivers for the same regional sex differences in mice^41^. These observations suggest that the basic topography of sex differences in human brain anatomy is laid down by genetic and environmental factors, which occur before and are largely independent of puberty. For some of these factors, like testicular hormones and sex chromosome dosage, we provide orthogonal support for their causal roles from our findings in rare disorders. It will be important for future work to also test for potential contributions of gender-based exposures^22–26^ to differences in subcortical anatomy between males and females. Our findings help guide this work by suggesting that gender effects occurring during or after puberty are not required for establishing the topography of sex differences in subcortical anatomy. Better delineating how sex and gender combine to influence the developing human brain is important for the accurate, comprehensive, and equitable advancement of knowledge in basic neuroscience and the many brain-based disorders displaying prominent differences between males and females.

## Methods

### Participants

All analyses of the datasets and the research protocol were approved by the National Institute of Mental Health Combined Neuroscience Institutional Review Board.

We computed sex differences in early adulthood using the Human Connectome Project Young Adults (HCP-YA) dataset, quantified out-of-sample reproducibility in middle adulthood using the mid-adult subset of the UK Biobank (UKB-MA) dataset, and defined their status in pre/early puberty using the 9-year-olds from the Adolescent Brain Cognitive Development (ABCD-9Y) dataset. The HCP structural MRI scans from 1,085 healthy young adults (592 females, 22-36y; 493 males, 22-37y) were downloaded from the HCP 1200 release (see ^50^ for details on recruitment procedures). The UKB structural MRI dataset included 669 participants (375 females, 45-50y; 294 males, 46-50y) within 5-year age range (45-50y) closest to that of the HCP sample (see ^51^ for details on recruitment procedures). The ABCD-9Y structural MRI dataset included 1,132 participants (569 females, 9-9.5y; 563 males, 9-9.5y) downloaded from Release 2.0.0 (see ^90^ for recruitment procedures).

Sex chromosome dosage effects were computed using a structural MRI dataset including 272 participants from two age-matched (two sample t-test p > 0.05) case-control sex chromosome aneuploidy (SCA) cohorts: XXY (n = 99, 6-25y; 92 control participants, 5-24y); and XYY (n = 34, 7-25y; 47 control participants, 5-24y). All individuals with SCA were reconfirmed to show a non-mosaic XXY or XYY karyotype by metaphase spread (or inspection of existing community records evidencing this in the 10 carriers unable to give blood). Participants were recruited through parent support organizations, the National Institute of Mental Health website and the National Institutes of Health Healthy Volunteers office. All participants had normal radiological findings and no history of brain injuries. All control participants were screened using a structured interview to ensure no history of neurodevelopmental or psychiatric disorders. All participants gave consent (or assent with parental consent as appropriate), and all protocols were completed at the NIH Clinical Center in Bethesda, Maryland.

The IGD dataset included 41 participants from an age-matched (two sample t-test p > 0.05) case-control cohort (19 male IGD patients, 14-41y; 22 male control patients, 13-42y). They were recruited through referrals from collaborators and local physicians or self-referrals with advertisement of the study on the NIH and Eunice Kennedy Shriver National Institute of Child Health and Human Development websites as well as ClinicalTrials.gov. Participants had a confirmed diagnosis of IGD with failure to undergo normal, age-appropriate pubertal development and low testosterone in the setting of low or inappropriately normal luteinizing hormone and follicle-stimulating hormone levels. Potential participants with additional pituitary hormone deficiencies or taking medications known to cause acquired hypogonadotropic hypogonadism (e.g., corticosteroids, chronic opioids) were excluded. All participants gave consent (or assent with parental consent as appropriate), and all protocols were completed at the National Institutes of Health Clinical Center in Bethesda, Maryland.

**Supplementary Table 1** provides demographic characteristics of normative (HCP-YA, UKB-MA, and ABCD-9Y) and clinical (XXY, XYY, and IGD) datasets.

### MRI Acquisition

T1-weighted (T1w) 3D magnetization-prepared rapid acquisition gradient-echo (MPRAGE) scans were acquired for all participants. The HCP-YA participants were scanned using 3T ConnectomeScanner (adapted from Siemens Skyra, Siemens Healthineers) with the following parameters: 0.7 mm isotropic resolution, FOV = 224 mm, TR = 2400 ms, TE = 2.14 ms, TI = 1000 ms, FA = 8°, bandwidth = 210 Hz per pixel, and GRAPPA = 2 (see ^91^ for details). The UKB-MA participants were scanned using 3T Siemens Skyra with the following parameters: 1 mm isotropic resolution, FOV = 256 mm, TR = 2000 ms, TE = 2.01 ms, TI = 880 ms, FA = 8°, bandwidth = 240 Hz per pixel, echo spacing = 6.1 ms and GRAPPA = 2 (see ^92^ for details). The ABCD-9Y participants were scanned using 3T scanners across different sites and platforms (including Siemens, Philips, or General Electric) with varying T1w sequences set at 1 mm isotropic resolution (see https://abcdstudy.org/scientists/protocols-mri/ for details). The SCA participants were scanned using 3T MR750 scanner (General Electric) with the following parameters: 1 mm isotropic resolution, FOV = 256 mm, TR = 2530 ms, TE = 3.5ms, TI = 1100 ms, FA = 7°, 195.3 Hz per pixel and GRAPPA = 2. The IGD participants were scanned using 3T MR750 scanner (General Electric) with the following parameters: 1 mm isotropic resolution, FOV = 256 mm, TR = 3205 ms, TE1-4 = 1.888, 3.832, 5.776, 7.72 ms, TI = 1150 ms, FA = 7°, bandwidth = 651 Hz per pixel, and GRAPPA = 2.

### MRI Data Processing

We used the MAGeT-Brain pipeline^48,49^ (**Figure 1A, left**) on T1w MRI scans to segment five subcortical structures (striatum, pallidum, thalamus, hippocampus, and amygdala) for measurement of bulk volume (**Figure 1A, middle**) and regional shape (**Figure 1A, right**). In brief, MAGeT-Brain segments subcortical structures by utilizing a small subset of high-quality, manually segmented atlases rather than a single atlas, reducing potential biases. It performs non-linear registration of these atlases to a representative subset of subjects, followed by propagation to all subjects to generate multiple candidate segmentations. These are then combined using a weighted voting strategy to define final anatomical labels. Surface models are subsequently defined by warping the surface mesh defined in the atlases to the space of individual subjects using the combined chain of nonlinear atlases-to-templates and templates-to-subjects transformations to model the surface representations of the structures. We then performed quality control based on visual inspection and histogram-based assessments of segmented volumes (see **Sup Table 2** for sample size per structure after quality control).

For a supplemental analysis of regional cortical anatomy, we used the automated recon-all pipeline of FreeSurfer (version 7.1.0) to process T1w MRI scans of the participants from the HCP-YA and ABCD-9Y datasets. This pipeline generates cortical surfaces from which morphometric features (i.e., cortical volume [CV], surface area [SA], and cortical thickness [CT]) were measured (as previously described^19^) for 360 cortical parcels in the multimodally derived HCP regional parcellation^93^. To control for overall brain size, we computed global CV and SA (by summation over 360 regions) and area-weighted mean CT (by averaging over 360 regions weighted by their surface area) for use in statistical models as a covariate. To control for variation in image and surface reconstruction quality, all included scans were verified as having Euler numbers (an index of topological defects in cortical surface reconstruction) greater than the recommended cutoff of −217^94^, and Euler number was included as a covariate in all statistical models estimating group differences in neuroanatomy.

### Statistical Analyses

#### Estimating Sex Differences in Subcortical Bulk Volumes and Regional Shape

Using MATLAB (version 2023a), we computed the standardized effects of sex differences (females vs. males) within the HCP-YA, UKB-MA, and ABCD-9Y datasets. For the HCP-YA dataset, we used the linear model of the form: Anatomy ∼ Sex + Age + Global Phenotype. Sex is defined as 1 for females (XX) and 0 for males (XY). Anatomy is bilateral bulk volume or vertex-wise surface area, and Global Phenotype is total brain tissue volume (for bulk volume analysis) or bilateral total surface area (for vertex-wise analysis). Age is age at scanning. For the UKB-MA and ABCD-9Y datasets, Site was added as a covariate to account for various research sites in which the participants were scanned. For the analyses of regional cortical anatomy, Anatomy is regional cortical volume (CV), surface area (SA), or cortical thickness (CT), while Global Phenotype is global CV, global SA, and area-weighted global mean CT. Euler was added as a covariate to control for surface reconstruction quality. All dependent variables were z-normalized across individuals, and numerical predictors were mean-centered across individuals, so that the beta coefficient for Sex represents a standardized effect size. For the supplemental analyses comparing different correction methods for overall brain size for estimating sex differences in subcortical shape from the HCP-YA dataset, “no correction” approach used the linear model of the form: Anatomy ∼ Sex + Age; “linear covariation” used the linear model of the form: Anatomy ∼ Sex + Age + Global Phenotype; and “allometric correction” used the linear model of the form: log_10_(Anatomy) ∼ Sex + Age + log_10_(Global Phenotype).

#### Defining X and Y Dosage Effects on Subcortical Bulk Volume and Regional Shape

Case-control differences (XXY vs. XY and XYY vs. XY) quantified X and Y dosage effects (respectively) within the SCA dataset. We applied a linear model for each case-control cohort defined as: Anatomy ∼ Group + Age + Global Phenotype, where Group is defined as 1 for XXY or XYY and 0 for XY. Anatomy is bilateral bulk volume or vertex-wise surface area, and Global Phenotype is total brain tissue volume (for bulk volume analysis) or bilateral total surface area (for vertex-wise analysis). Age is age at scanning. All dependent variables were z-normalized, and numeric predictor variables were mean-centered across individuals so that the beta coefficient associated with Group represents a standardized effect size for a unit increase in X- and Y-chromosome dosage.

#### Defining the Effects of Intact Testicular Hormone Production on Subcortical Bulk Volume and Regional Shape

We quantified the effects of intact testicular hormone production (typically developing XY controls vs. IGD XY individuals) using a linear model of the form: Anatomy ∼ Group + Age + Global Phenotype, where Group is defined as 1 for typical XY males and 0 for IGD male patients. Anatomy is bilateral bulk volume or vertex-wise surface area, and Global Phenotype is total brain tissue volume (for bulk volume analysis) or bilateral total surface area (for vertex-wise analysis). Age is age at scanning. All dependent variables were z-normalized across individuals, and numeric predictors were mean-centered across individuals, so that the beta coefficient associated with Group represents the standardized effect size of intact testicular hormone production.

#### Controlling for Multiple Comparisons

For bulk level analyses of normative sex differences, sex chromosome dosage, and testicular hormone production, we used Bonferroni correction across the 5 subcortical structures to control for multiple comparisons (p_BF_ = 5*p) to be thresholded at p_BF_ < 0.05. For regional shape analyses, false discovery rate (FDR) correction with p_FDR_ < 0.05 accounted for multiple comparisons across subcortical vertices in both hemispheres within each structure. For the analyses of regional cortical anatomy, FDR with p_FDR_ < 0.05 accounted for multiple comparisons across 360 cortical regions of the HCP multimodal parcellation.

#### Constructing Null Spatial Models for Comparing Two Maps of Regional Shape

The statistical significance of observed spatial correlations between two maps for regional shape was tested against a spatial null model that accounts for potential false positives arising from the spatial autocorrelation inherent in the neuroimaging phenotypes. The spatial null model was constructed based on 10,000 Moran spectral randomizations^52,53^ of one of the two maps being compared. In brief, Moran spectral randomization generates spatially constrained null brain maps by decomposing the source map into spatial eigenvectors and randomly perturbing their coefficients, producing surrogate data that preserves the source map’s spatial autocorrelation. The percentile position of the spatial correlation between the two original maps within this null distribution determined statistical significance at p_MSR_ < 0.05, followed by Bonferroni correction across the 5 subcortical structures at p_MSR-BF_ = 5*p_MSR_ to be thresholded at p_MSR-BF_ < 0.05. For the analysis of cortical sex differences, we applied a spin-based spatial permutation test^95^ followed by Bonferroni correction across 3 cortical features (p_SPIN-BF_ = 3*p_SPIN_) to be thresholded at p_SPIN-BF_ < 0.05. Note that while a spin-based spatial permutation test exists for striatum, pallidum, and thalamus^96^, it is currently unavailable for hippocampus and amygdala.

### Reproducibility of Sex Differences within Early Adulthood as a Function of Sample Size

We quantified the within-sample reproducibility of sex differences in subcortical shape using 10,000 instances of split-half, sex-balanced random subsampling of the HCP-YA dataset at varying sample sizes (from 50 to 450, with increments of 50) and without replacement. For example, in each repetition and at the split-half size of 450, sex differences were recomputed (using the approach described in “*Estimating Sex Differences in Subcortical Bulk Volumes and Regional Shape*” above) in a split-half sample of 450 (225 females and 225 males) and the other split-half sample of 450 (225 females and 225 males), and the Pearson correlation between the two resulting standardized effect size maps (not thresholded) was computed. This procedure generated a distribution of 10,000 spatial correlations for each sample size, thereby offering an objective measure of the within-sample reproducibility of sex differences in subcortical shape and its relation to sample size.

### Reproducibility of Sex Differences between Early and Middle Adulthood

To assess the out-of-sample replicability of sex differences in subcortical shape of each structure, we computed the vertex-wise Pearson correlations between the sex differences from HCP-YA and UKB-MA datasets. The statistical significance of these correlations was determined based on 10,000 Moran spectral randomizations of HCP-YA sex differences, followed by Bonferroni correction at p_MSR-BF_ < 0.05 across the 5 subcortical structures. Additionally, we computed the regression slope predicting UKB-MA sex differences as the dependent variable using the HCP-YA sex differences as the predictor for each structure. Consequently, a slope of >1 indicates that the sex differences in subcortical shape across a given structure tend to be stronger in UKB-MA vs. HCP-YA, while a slope of <1 indicates that UKB-MA sex effects tend to be weaker than in HCP-YA.

To formally compare the regional congruence of statistically significant sex differences between the HCP-YA and UKB-MA datasets, we generated a conjunction map defining 3 levels of agreement (HCP-YA and UKB-MA both showing a statistically significant sex difference in the same direction; HCP-YA and UKB-MA both showing the same direction of sex difference, but only reaching statistical significance in one dataset; HCP-YA and UKB-MA both showing the same direction of statistically non-significant sex difference) and 3 levels of disagreement (HCP-YA and UKB-MA showing a statistically significant sex difference in opposite directions; HCP-YA and UKB-MA showing a sex difference in opposite directions which reached statistical significance in one dataset; HCP-YA and UKB-MA showing statistically non-significant sex differences in opposite directions). To quantify the extent to which sex-biased subcortical shape agrees between the two datasets, we calculated the proportion of subcortical surface that fell into each of these possible conjunction agreement/disagreement cases. To determine the statistical significance of regional agreement between the two datasets at the structure-level, we compared the combined areal proportion of the subcortical surface in agreement (in all 3 levels) to the null distribution of agreement proportions after 10,000 Moran spectral randomizations of HCP-YA sex difference maps, followed by Bonferroni correction across the 5 subcortical structures at p_MSR-BF_ < 0.05.

### Consistency of Sex Differences between Early Adulthood and Pre/Early Puberty

To assess the consistency of sex differences in subcortical shape of each structure, we computed the vertex-wise Pearson correlations between the sex differences from HCP-YA and ABCD-9Y datasets. The statistical significance of these correlations was determined based on 10,000 Moran spectral randomizations of HCP-YA sex differences, followed by Bonferroni correction at p_MSR-BF_ < 0.05 across the 5 subcortical structures. Additionally, we computed the regression slope predicting HCP-YA sex differences as the dependent variable using the ABCD-9Y sex differences as the predictor for each structure. Consequently, a slope of >1 indicates that the sex differences in subcortical shape across a given structure tend to be stronger in HCP-YA vs. ABCD-9Y, which is compatible with the notion that sex effects have accentuated from pre/early puberty to early adulthood. Conversely, a slope of <1 indicates that HCP-YA sex effects tend to be weaker than in ABCD-9Y, suggesting attenuation from pre/early puberty to early adulthood.

We then assessed the consistency of the statistical significance and direction of sex differences in subcortical shape between pre/early puberty and early adulthood in a region-by-region manner. To formally examine the regional congruence between these two cohorts, we partitioned the sex differences in early adulthood by their status in pre/early puberty with the following categories: never sex-biased (sex-biases are non-significant at both pre/early puberty and early adulthood); established (female- and male-biases are significant in the same direction in pre/early puberty and early adulthood); emerging (female- and male-biases are significant in early adulthood but non-significant at pre/early puberty); diminishing (female- and male-biases are significant in pre/early puberty but non-significant at early adulthood); and inverted (female- and male-biases are significantly in the opposite directions between pre/early puberty and early adulthood). To determine the statistical significance of consistency between the two datasets, we compared the combined areal proportion of the subcortical surface in consistency (falling into established, emerging, or never sex-biased) to the null distribution of consistent proportions after 10,000 Moran spectral randomizations of HCP-YA sex difference maps, followed by Bonferroni correction across the 5 subcortical structures at p_MSR-BF_ < 0.05. In parallel, we assessed the consistency for sex differences in regional cortical anatomy between pre/early puberty and early adulthood following identical procedures, as an extension to our previous study focusing on cortical sex differences^19^.

### Relationship of Sex Differences in Subcortical Shape with X/Y Dosage and Testicular Hormone Production

We annotated regions of statistically significant sex differences in subcortical shape by the regional congruence of X (XXY vs. XY), Y (XYY vs. XY), and T (XY controls vs. IGD XY individuals) effects. Specifically, we categorized all subcortical vertices according to whether the direction of sex differences was congruent with the significance and direction of X, Y, and T effects. Here, we defined congruent effects as those where the direction of X, Y, and T effects is compatible with the direction of normative sex differences (i.e., female-biased: XXY>XY, XYY<XY, or control<IGD; male-biased: XXY<XY, XYY>XY, or control>IGD). In these analyses, we differentiated between “hard” congruences (where X, Y, and T effects reached statistical significance) and “soft” congruences (where X, Y, and T effects fell below statistical significance). To assess congruence at the structure-level, these proportions were then compared to the null distribution of the proportions computed after 10,000 Moran spectral randomizations of sex difference maps to determine statistical significance (p_MSR_ < 0.05), followed by Bonferroni correction (p_MSR-BF_ < 0.05) across the 5 subcortical structures to control for multiple comparisons.

## Data availability

Group-level statistical parameters (standardized effect sizes, nominal p values and p values adjusted for multiple comparisons) for the effects of sex, sex chromosome dosage, and testicular hormone production will be provided as source data upon publication. The HCP S1200 data release (March 2017) is publicly available at https://db.humanconnectome.org/. UK Biobank data (2020 release) are available via application at https://www.ukbiobank.ac.uk. ABCD data (Release 2.0.0) are available via the NIMH Data Archive (https://nda.nih.gov/). For the HCP, UKB, and ABCD datasets, the data access agreements and privacy policies prohibit sharing participant-level data such as neuroimaging-derived variables. Data from the NIH clinical cohorts (SCAs and IGD) are not publicly available due to participant privacy, but are available from the corresponding author upon reasonable request and completion of a data use agreement, consistent with NIH IRB protocols.

## Code availability

Subcortical segmentation and shape analysis used the publicly available MAGeT-Brain pipeline (https://github.com/CoBrALab/MAGeTbrain). Analysis code is publicly available at https://github.com/TheHyoLee/sex_differences_in_human_subcortical_anatomy.

## Acknowledgments

We thank all study participants for taking part in this research. This research was supported by the Intramural Research Program of the NIMH (1ZIAMH002949-10).

## Author Contribution Statement

H.M.L. and A.R. conceptualized the study. H.M.L. developed the methodology, performed the formal analysis, and wrote the original draft. L.C., E.G., D.M., A.T., and Angela D. provided essential resources, including data from the Human Connectome Project, UK Biobank, and clinical IGD cohort. G.A.D. and M.M.C. provided the essential data processing pipeline (MAGeT-Brain) and support. F.M.L. and L.C. curated the clinical neuroimaging datasets. A.R. supervised the research, secured funding and drafted the manuscript. All authors (H.M.L., S.L., F.M.L., L.C., E.G., Alex D., D.M., A.T., J.P.L., G.A.D., M.M.C., K.F.B., Angela D., and A.R.) reviewed and edited the manuscript.

## Competing Interests Statement

The authors declare no competing interests.

## Notes

### Competing Interest Statement

The authors have declared no competing interest.

### Summary of Updates

A typo in Discussion and another in Results are corrected.

## References

1 Eliot, L., Ahmed, A., Khan, H. & Patel, J. Dump the “dimorphism”: Comprehensive synthesis of human brain studies reveals few male-female differences beyond size. Neuroscience & Biobehavioral Reviews 125, 667–697 (2021). 10.1016/j.neubiorev.2021.02.026

2 DeCasien, A. R., Guma, E., Liu, S. & Raznahan, A. Sex differences in the human brain: a roadmap for more careful analysis and interpretation of a biological reality. Biology of Sex Differences 13, 43 (2022). 10.1186/s13293-022-00448-w

3 GBD 2019 Mental Disorders Collaborators. Global, regional, and national burden of 12 mental disorders in 204 countries and territories, 1990–2019: a systematic analysis for the Global Burden of Disease Study 2019. The Lancet Psychiatry 9, 137–150 (2022). 10.1016/S2215-0366(21)00395-3

4 Steinmetz, J. D. et al. Global, regional, and national burden of disorders affecting the nervous system, 1990–2021: a systematic analysis for the Global Burden of Disease Study 2021. The Lancet Neurology 23, 344–381 (2024). 10.1016/S1474-4422(24)00038-3

5 Duerden, E. G., Chakravarty, M. M., Lerch, J. P. & Taylor, M. J. Sex-Based Differences in Cortical and Subcortical Development in 436 Individuals Aged 4–54 Years. Cerebral Cortex 30, 2854–2866 (2020). 10.1093/cercor/bhz279

6 Kalantar-Hormozi, H. et al. A cross-sectional and longitudinal study of human brain development: The integration of cortical thickness, surface area, gyrification index, and cortical curvature into a unified analytical framework. NeuroImage 268, 119885 (2023). 10.1016/j.neuroimage.2023.119885

7 Küchenhoff, S. et al. Relating sex-bias in human cortical and hippocampal microstructure to sex hormones. Nature Communications 15, 7279 (2024). 10.1038/s41467-024-51459-7

8 Lenroot, R. K. & Giedd, J. N. Brain development in children and adolescents: insights from anatomical magnetic resonance imaging. Neuroscience & biobehavioral reviews 30, 718–729 (2006).

9 Sanchis-Segura, C. et al. Sex differences in gray matter volume: how many and how large are they really? Biology of Sex Differences 10, 32 (2019). 10.1186/s13293-019-0245-7

10 Williams, C. M., Peyre, H., Toro, R. & Ramus, F. Neuroanatomical norms in the UK Biobank: The impact of allometric scaling, sex, and age. Human Brain Mapping 42, 4623–4642 (2021). 10.1002/hbm.25572

11 Ritchie, S. J. et al. Sex Differences in the Adult Human Brain: Evidence from 5216 UK Biobank Participants. Cerebral Cortex 28, 2959–2975 (2018). 10.1093/cercor/bhy109

12 Liu, S., Seidlitz, J., Blumenthal, J. D., Clasen, L. S. & Raznahan, A. Integrative structural, functional, and transcriptomic analyses of sex-biased brain organization in humans. Proceedings of the National Academy of Sciences 117, 18788–18798 (2020). 10.1073/pnas.1919091117

13 Ruigrok, A. N. V. et al. A meta-analysis of sex differences in human brain structure. Neuroscience & Biobehavioral Reviews 39, 34–50 (2014). 10.1016/j.neubiorev.2013.12.004

14 Giedd, J. N., Raznahan, A., Mills, K. L. & Lenroot, R. K. Review: magnetic resonance imaging of male/female differences in human adolescent brain anatomy. Biology of Sex Differences 3, 19 (2012). 10.1186/2042-6410-3-19

15 Raznahan, A. et al. How Does Your Cortex Grow? The Journal of Neuroscience 31, 7174–7177 (2011). 10.1523/jneurosci.0054-11.2011

16 Luders, E. & Toga, A. W. in Progress in Brain Research Vol. 186 (ed Ivanka Savic) 2–12 (Elsevier, 2010).

17 Zhou, X. et al. Choice of Voxel-based Morphometry processing pipeline drives variability in the location of neuroanatomical brain markers. Communications Biology 5, 913 (2022). 10.1038/s42003-022-03880-1

18 Williams, C. M., Peyre, H., Toro, R. & Ramus, F. Sex differences in the brain are not reduced to differences in body size. Neuroscience & Biobehavioral Reviews 130, 509–511 (2021). 10.1016/j.neubiorev.2021.09.015

19 Lee, H. M. et al. Regional sex differences in human cortical anatomy vary in their morphometric bases and overlap with sex chromosomal and gonadal influences. Nature Communications 17, 7503 (2026). 10.1038/s41467-026-74274-8

20 Reardon, P. K. et al. An Allometric Analysis of Sex and Sex Chromosome Dosage Effects on Subcortical Anatomy in Humans. The Journal of Neuroscience 36, 2438–2448 (2016). 10.1523/jneurosci.3195-15.2016

21 Vijayakumar, N. et al. The effects of puberty and its hormones on subcortical brain development. Comprehensive Psychoneuroendocrinology 7, 100074 (2021). 10.1016/j.cpnec.2021.100074

22 Portengen, C. M., van Baar, A. L. & Endendijk, J. J. A neurocognitive approach to studying processes underlying parents’ gender socialization. Frontiers in Psychology 13, 1054886 (2023).

23 Wu, H. et al. Occupational neuroplasticity in the human brain: a critical review and meta-analysis of neuroimaging studies. Frontiers in human neuroscience 14, 215 (2020).

24 Graves, B. S., Hall, M. E., Dias-Karch, C., Haischer, M. H. & Apter, C. Gender differences in perceived stress and coping among college students. PLOS ONE 16, e0255634 (2021). 10.1371/journal.pone.0255634

25 Forbes, C. E. & Leitner, J. B. Stereotype threat engenders neural attentional bias toward negative feedback to undermine performance. Biological Psychology 102, 98–107 (2014). 10.1016/j.biopsycho.2014.07.007

26 Zugman, A. et al. Country-level gender inequality is associated with structural differences in the brains of women and men. Proceedings of the National Academy of Sciences 120, e2218782120 (2023). 10.1073/pnas.2218782120

27 De Vries, G. J. et al. A Model System for Study of Sex Chromosome Effects on Sexually Dimorphic Neural and Behavioral Traits. The Journal of Neuroscience 22, 9005–9014 (2002). 10.1523/jneurosci.22-20-09005.2002

28 Cruz, K. G. et al. Cortical-subcortical interactions in goal-directed behavior. Physiological Reviews 103, 347–389 (2022). 10.1152/physrev.00048.2021

29 Miletić, S. et al. Joint models reveal human subcortical underpinnings of choice and learning behavior. Proceedings of the National Academy of Sciences 122, e2502269122 (2025). 10.1073/pnas.2502269122

30 Nazzi, A., Favaretto, C., Vallesi, A., Corbetta, M. & Allegra, M. Switching Patterns of Cortical–Subcortical Interaction in the Human Brain. The Journal of Neuroscience 45, e1855242025 (2025). 10.1523/JNEUROSCI.1855-24.2025

31 Gupta, R., Koscik, T. R., Bechara, A. & Tranel, D. The amygdala and decision-making. Neuropsychologia 49, 760–766 (2011). 10.1016/j.neuropsychologia.2010.09.029

32 Courellis, H. S. et al. Abstract representations emerge in human hippocampal neurons during inference. Nature 632, 841–849 (2024). 10.1038/s41586-024-07799-x

33 Frank, D. et al. Human hippocampal ripples tune cortical responses based on predicted uncertainty. Nature Neuroscience 29, 1987–1998 (2026). 10.1038/s41593-026-02345-6

34 Pauli, W. M., O’Reilly, R. C., Yarkoni, T. & Wager, T. D. Regional specialization within the human striatum for diverse psychological functions. Proceedings of the National Academy of Sciences 113, 1907–1912 (2016). 10.1073/pnas.1507610113

35 Griffiths, B. J. et al. Rhythmic interactions between the mediodorsal thalamus and prefrontal cortex precede human visual perception. Nature Communications 13, 3736 (2022). 10.1038/s41467-022-31407-z

36 Bruno, R. M. & Sakmann, B. Cortex Is Driven by Weak but Synchronously Active Thalamocortical Synapses. Science 312, 1622–1627 (2006). 10.1126/science.1124593

37 Forger, N. G. et al. Deletion of Bax eliminates sex differences in the mouse forebrain. Proceedings of the National Academy of Sciences 101, 13666–13671 (2004). 10.1073/pnas.0404644101

38 Campi, K. L., Jameson, C. E. & Trainor, B. C. Sexual dimorphism in the brain of the monogamous California mouse (Peromyscus californicus). Brain, behavior and evolution 81, 236–249 (2013).

39 Qiu, L. R. et al. Mouse MRI shows brain areas relatively larger in males emerge before those larger in females. Nature Communications 9, 2615 (2018). 10.1038/s41467-018-04921-2

40 Guma, E. et al. Comparative neuroimaging of sex differences in human and mouse brain anatomy. eLife 13, RP92200 (2024). 10.7554/eLife.92200

41 Lerch, J. P. et al. Gonadal and sex chromosomal contributions to sex differences in mammalian brain organization. bioRxiv [Preprint], version 1 (2026). 10.64898/2026.01.10.698782

42 Gorski, R. A., Gordon, J. H., Shryne, J. E. & Southam, A. M. Evidence for a morphological sex difference within the medial preoptic area of the rat brain. Brain Research 148, 333–346 (1978). 10.1016/0006-8993(78)90723-0

43 Bleier, R., Byne, W. & Siggelkow, I. Cytoarchitectonic sexual dimorphisms of the medial preoptic and anterior hypothalamic areas in guinea pig, rat, hamster, and mouse. Journal of Comparative Neurology 212, 118–130 (1982). 10.1002/cne.902120203

44 Vousden, D. A. et al. Impact of X/Y genes and sex hormones on mouse neuroanatomy. NeuroImage 173, 551–563 (2018). 10.1016/j.neuroimage.2018.02.051

45 Corre, C. et al. Separate effects of sex hormones and sex chromosomes on brain structure and function revealed by high-resolution magnetic resonance imaging and spatial navigation assessment of the Four Core Genotype mouse model. Brain Structure and Function 221, 997–1016 (2016). 10.1007/s00429-014-0952-0

46 Arnold, A. P. A general theory of sexual differentiation. Journal of Neuroscience Research 95, 291–300 (2017). 10.1002/jnr.23884

47 Fish, A. M. et al. Sex-biased trajectories of amygdalo-hippocampal morphology change over human development. NeuroImage 204, 116122 (2020). 10.1016/j.neuroimage.2019.116122

48 Pipitone, J. et al. Multi-atlas segmentation of the whole hippocampus and subfields using multiple automatically generated templates. NeuroImage 101, 494–512 (2014).

49 Chakravarty, M. M. et al. Performing label-fusion-based segmentation using multiple automatically generated templates. Human Brain Mapping 34, 2635–2654 (2013). 10.1002/hbm.22092

50 Van Essen, D. C. et al. The WU-Minn Human Connectome Project: An overview. NeuroImage 80, 62–79 (2013). 10.1016/j.neuroimage.2013.05.041

51 Sudlow, C. et al. UK Biobank: An Open Access Resource for Identifying the Causes of a Wide Range of Complex Diseases of Middle and Old Age. PLOS Medicine 12, e1001779 (2015). 10.1371/journal.pmed.1001779

52 Wagner, H. H. & Dray, S. Generating spatially constrained null models for irregularly spaced data using Moran spectral randomization methods. Methods in Ecology and Evolution 6, 1169–1178 (2015).

53 Markello, R. D. et al. neuromaps: structural and functional interpretation of brain maps. Nature Methods (2022). 10.1038/s41592-022-01625-w

54 Nielsen, J. & Wohlert, M. Chromosome abnormalities found among 34910 newborn children: results from a 13-year incidence study in Århus, Denmark. Human Genetics 87, 81–83 (1991). 10.1007/BF01213097

55 Nadig, A. et al. Carriage of Supernumerary Sex Chromosomes Decreases the Volume and Alters the Shape of Limbic Structures. eneuro 5, ENEURO.0265-0218.2018 (2018). 10.1523/eneuro.0265-18.2018

56 Seidlitz, J. et al. Transcriptomic and cellular decoding of regional brain vulnerability to neurogenetic disorders. Nature Communications 11, 3358 (2020). 10.1038/s41467-020-17051-5

57 Hong, D. S. et al. Influence of the X-Chromosome on Neuroanatomy: Evidence from Turner and Klinefelter Syndromes. The Journal of Neuroscience 34, 3509–3516 (2014). 10.1523/jneurosci.2790-13.2014

58 Laitinen, E.-M. et al. Incidence, phenotypic features and molecular genetics of Kallmann syndrome in Finland. Orphanet journal of rare diseases 6, 1–10 (2011).

59 Dodé, C. & Hardelin, J.-P. Kallmann syndrome. European Journal of Human Genetics 17, 139–146 (2009). 10.1038/ejhg.2008.206

60 Kim, S.-H. Congenital hypogonadotropic hypogonadism and Kallmann syndrome: past, present, and future. Endocrinology and metabolism 30, 456 (2015).

61 Rodprasert, W., Virtanen, H. E., Mäkelä, J.-A. & Toppari, J. Hypogonadism and Cryptorchidism. Frontiers in Endocrinology 10, 906 (2020). 10.3389/fendo.2019.00906

62 Cabrejas Gómez, M. d. C., Vicente Vicente, M. Á., Antón Miguel, M. Á. & Urcelay Rojo, M. Late-diagnosed Kallmann syndrome. Endocrinología y Nutrición (English Edition) 62, 106–108 (2015). 10.1016/j.endoen.2014.10.002

63 Jallepalli, D. & Dang, S. Cortico-subcortical converging organization at rest. Scientific Reports 15, 32133 (2025). 10.1038/s41598-025-18023-9

64 Prakash, N., Abu Irqeba, A. & Corbin, J. G. Development and function of the medial amygdala. Trends in Neurosciences 48, 22–32 (2025). 10.1016/j.tins.2024.11.004

65 Torrisi, S. et al. Resting state connectivity of the bed nucleus of the stria terminalis at ultra-high field. Human Brain Mapping 36, 4076–4088 (2015). 10.1002/hbm.22899

66 Yeo, T. B. T. et al. The organization of the human cerebral cortex estimated by intrinsic functional connectivity. Journal of Neurophysiology 106, 1125–1165 (2011). 10.1152/jn.00338.2011

67 Liu, S. et al. Robust but independent sex differences in human brain function, structure, and behavior. Nature Communications 17, 6694 (2026). 10.1038/s41467-026-73262-2

68 Arnold, A. P. Sexual differentiation of brain and other tissues: Five questions for the next 50 years. Hormones and Behavior 120, 104691 (2020). 10.1016/j.yhbeh.2020.104691

69 Zugman, A., Jett, L., Antonacci, C., Winkler, A. M. & Pine, D. S. A systematic review and meta-analysis of resting-state fMRI in anxiety disorders: Need for data sharing to move the field forward. Journal of Anxiety Disorders 99, 102773 (2023). 10.1016/j.janxdis.2023.102773

70 Herting, M. M. et al. Correspondence Between Perceived Pubertal Development and Hormone Levels in 9-10 Year-Olds From the Adolescent Brain Cognitive Development Study. Frontiers in Endocrinology 11, 549928 (2021). 10.3389/fendo.2020.549928

71 Raznahan, A. et al. Globally Divergent but Locally Convergent X- and Y-Chromosome Influences on Cortical Development. Cerebral Cortex 26, 70–79 (2014). 10.1093/cercor/bhu174

72 Manara, R. et al. Brain Changes in Kallmann Syndrome. American Journal of Neuroradiology 35, 1700–1706 (2014). 10.3174/ajnr.A3946

73 Shirazi, T. N. et al. Low Perinatal Androgens Predict Recalled Childhood Gender Nonconformity in Men. Psychological Science 33, 343–353 (2022). 10.1177/09567976211036075

74 Mallard, T. T. et al. X-chromosome influences on neuroanatomical variation in humans. Nature Neuroscience 24, 1216–1224 (2021). 10.1038/s41593-021-00890-w

75 Cole, K. M. et al. The NIMH Intramural Longitudinal Study of the Endocrine and Neurobiological Events Accompanying Puberty: Protocol and rationale for methods and measures. NeuroImage 234, 117970 (2021). 10.1016/j.neuroimage.2021.117970

76 Heller, C. et al. Hormonal milieu influences whole-brain structural dynamics across the menstrual cycle using dense sampling in multiple individuals. Nature Neuroscience 28, 2588–2600 (2025).

77 Heller, C. et al. Menarche onset is an inflection point for mental health and brain development. bioRxiv [Preprint] (2026). 10.64898/2026.04.13.718198

78 Rizor, E. J. et al. Menstrual cycle-driven hormone concentrations co-fluctuate with white and gray matter architecture changes across the whole brain. Human Brain Mapping 45, e26785 (2024).

79 Pritschet, L. et al. Neuroanatomical changes observed over the course of a human pregnancy. Nature Neuroscience 27, 2253–2260 (2024). 10.1038/s41593-024-01741-0

80 Martínez-García, M., Jacobs, E. G., de Lange, A.-M. G. & Carmona, S. Advancing the neuroscience of human pregnancy. Nature Neuroscience 27, 805–807 (2024). 10.1038/s41593-024-01629-z

81 Mosconi, L. et al. Menopause impacts human brain structure, connectivity, energy metabolism, and amyloid-beta deposition. Scientific Reports 11, 10867 (2021). 10.1038/s41598-021-90084-y

82 Taylor, C. M., Pritschet, L., Yu, S. & Jacobs, E. G. Applying a Women’s Health Lens to the Study of the Aging Brain. Frontiers in Human Neuroscience 13, 224 (2019). 10.3389/fnhum.2019.00224

83 Anawalt, B. D. & Merriam, G. R. NEUROENDOCRINE AGING IN MEN: Andropause and Somatopause. Endocrinology and Metabolism Clinics of North America 30, 647–669 (2001). 10.1016/S0889-8529(05)70206-1

84 Ogoniak, L. et al. Role of genetics in the age-related testosterone decline in men: a UK Biobank study. European Journal of Endocrinology 193, 197–203 (2025). 10.1093/ejendo/lvaf143

85 Meaney, M. J. & McEwen, B. S. Testosterone implants into the amygdala during the neonatal period masculinize the social play of juvenile female rats. Brain Research 398, 324–328 (1986). 10.1016/0006-8993(86)91492-7

86 Arimatsu, Y. Short- and long-term influences of neonatal sex steroids on α-bungarotoxin binding capacity in the mouse amygdala. Neuroscience 9, 873–877 (1983). 10.1016/0306-4522(83)90275-0

87 Kraatz, G., Xie, H. T. H., Long, H. & Walker, C.-D. Neonatal estradiol and early adversity interact to modify basolateral amygdala morphology and adult behavior in female rats. Journal of Neuroendocrinology 37, e13483 (2025). 10.1111/jne.13483

88 De Lorme, K. C., Schulz, K. M., Salas-Ramirez, K. Y. & Sisk, C. L. Pubertal testosterone organizes regional volume and neuronal number within the medial amygdala of adult male Syrian hamsters. Brain Research 1460, 33–40 (2012). 10.1016/j.brainres.2012.04.035

89 Morris, J. A., Jordan, C. L., King, Z. A., Northcutt, K. V. & Breedlove, S. M. Sexual dimorphism and steroid responsiveness of the posterodorsal medial amygdala in adult mice. Brain Research 1190, 115–121 (2008). 10.1016/j.brainres.2007.11.005

90 Karcher, N. R. & Barch, D. M. The ABCD study: understanding the development of risk for mental and physical health outcomes. Neuropsychopharmacology 46, 131–142 (2021). 10.1038/s41386-020-0736-6

91 Van Essen, D. C. et al. The Human Connectome Project: A data acquisition perspective. NeuroImage 62, 2222–2231 (2012). 10.1016/j.neuroimage.2012.02.018

92 Miller, K. L. et al. Multimodal population brain imaging in the UK Biobank prospective epidemiological study. Nature Neuroscience 19, 1523–1536 (2016). 10.1038/nn.4393

93 Glasser, M. F. et al. A multi-modal parcellation of human cerebral cortex. Nature 536, 171–178 (2016).

94 Rosen, A. F. G. et al. Quantitative assessment of structural image quality. NeuroImage 169, 407–418 (2018). 10.1016/j.neuroimage.2017.12.059

95 Váša, F. et al. Adolescent Tuning of Association Cortex in Human Structural Brain Networks. Cerebral Cortex 28, 281–294 (2017). 10.1093/cercor/bhx249

96 Blostein, N. et al. Morphological and anatomical variations in subcortical anatomy between humans and chimpanzees associated with heritability patterns related to human behavioral traits. Communications Biology 9, 852 (2026). 10.1038/s42003-026-10066-6

