## Supplemental Figures for "Sex Differences in Human Subcortical Anatomy are Robust and Replicable in Adulthood, Already Evident Prepubertally, and Aligned with Sex Chromosomal and Gonadal Influences"

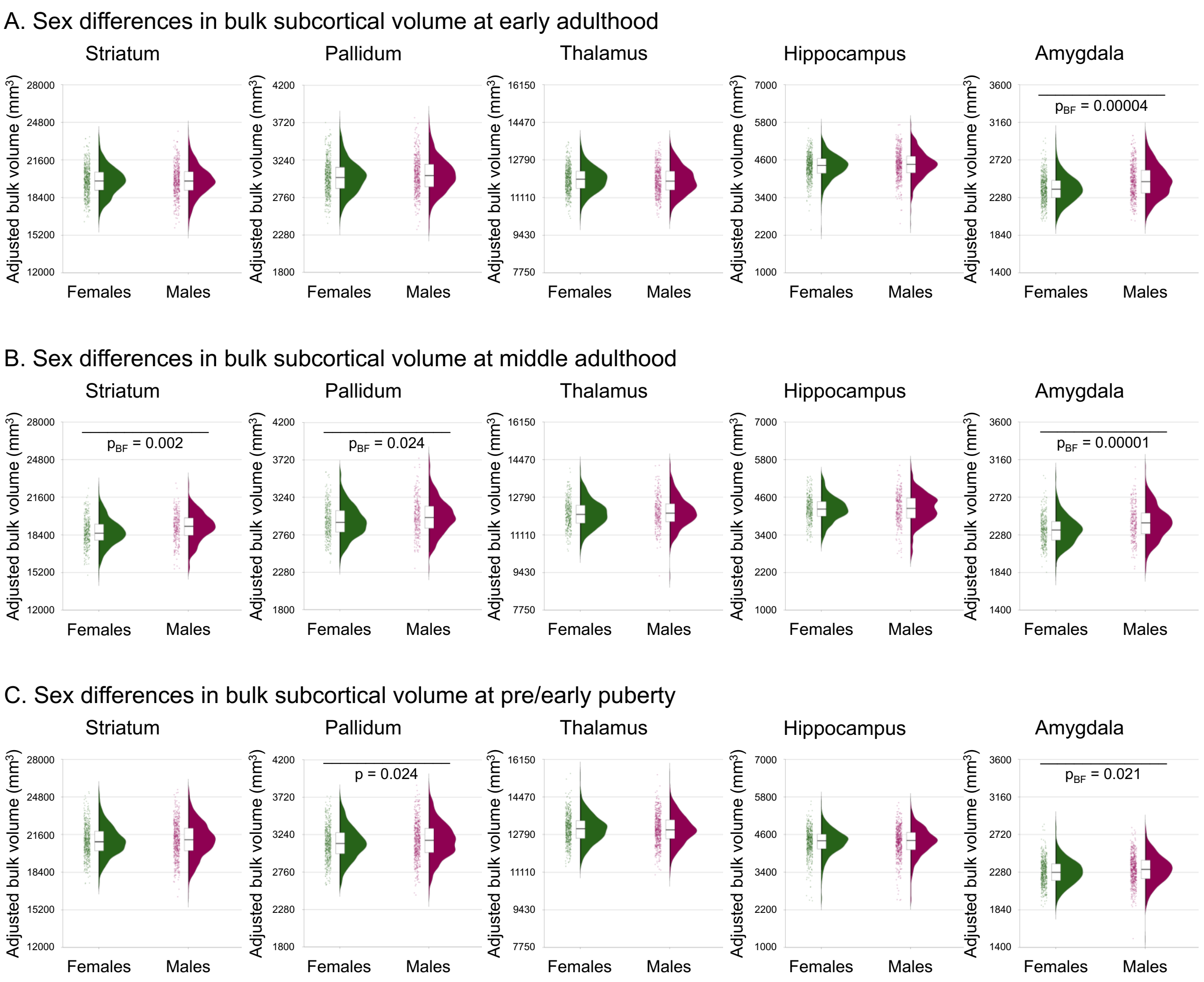


**Supplemental Figure 1. Sex differences in bulk subcortical volumes across early adulthood, middle adulthood, and pre/early puberty. A.** Bulk volume of subcortical structures (adjusted for age and total brain volume) for females (green) and males (magenta) in early adulthood (HCP-YA). **B.** Bulk volume of subcortical structures (adjusted for age, total brain volume, and site) for females (green) and males (magenta) in middle adulthood (UKB-MA). **C.** Bulk volume of subcortical structures (adjusted for age, total brain volume, and site) for females (green) and males (magenta) in pre/early puberty (ABCD-9Y). For all panels, the boxes, histograms, and jittered dots indicate median bounded by first and third quartiles, distribution, and individual values. Horizontal bars indicate structures showing significant sex differences at nominal $p < 0.05$ or Bonferroni-corrected $p_{BF} < 0.05$.

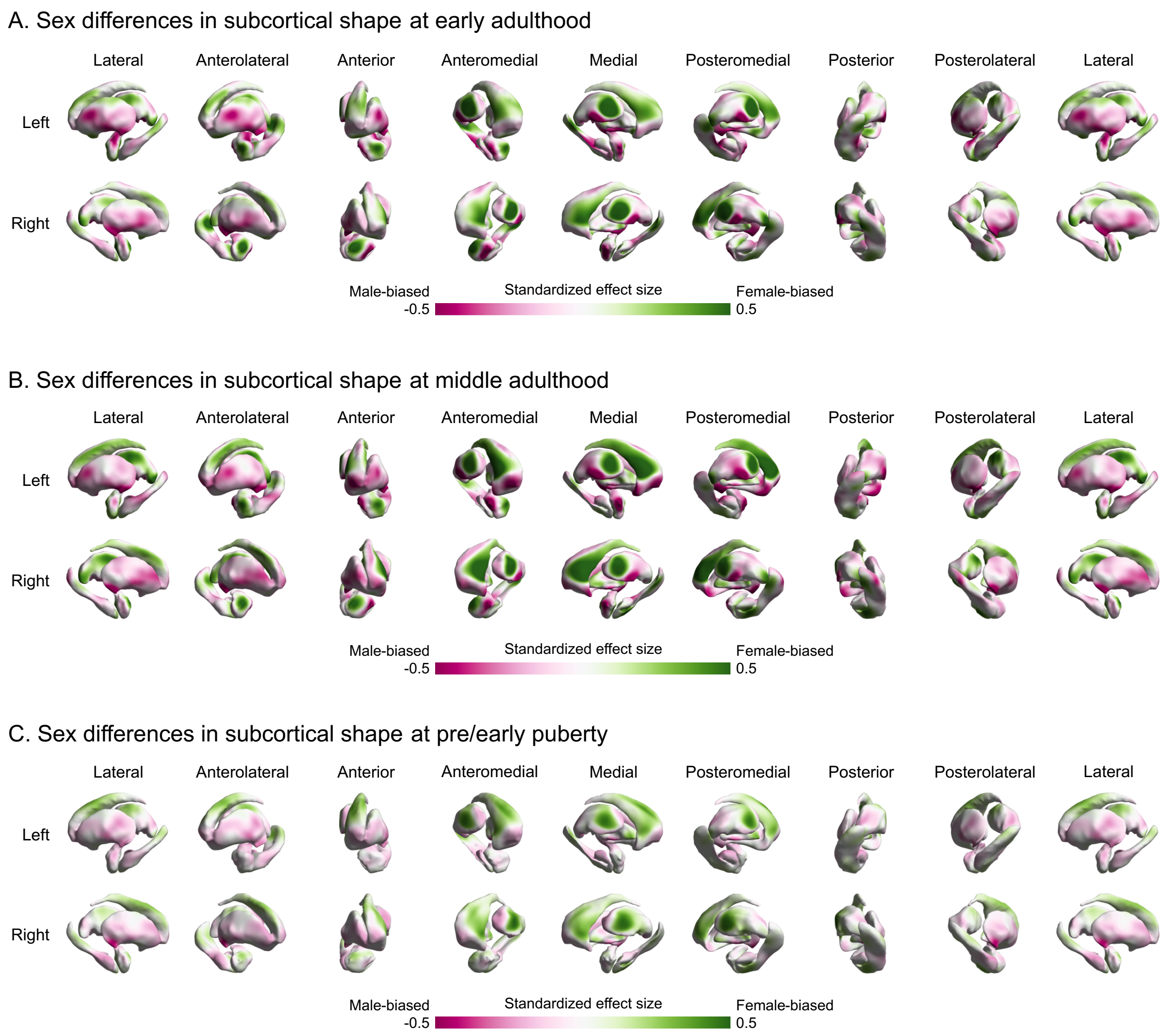


**Supplemental Figure 2. Sex differences in regional subcortical shape across early adulthood, middle adulthood, and pre/early puberty. A.** Standardized effect sizes for sex differences in regional surface area (controlling for age and total surface area) showing female-biases (green) and male-biases (magenta) in early adulthood (HCP-YA). **B.** Standardized effect sizes for sex differences in regional surface area (controlling for age, total surface area, and site) showing female-biases (green) and male-biases (magenta) in middle adulthood (UKB-MA). **C.** Standardized effect sizes for sex differences in regional surface area (controlling for age, total surface area, and site) showing female-biases (green) and male-biases (magenta) in pre/early puberty (ABCD-9Y).

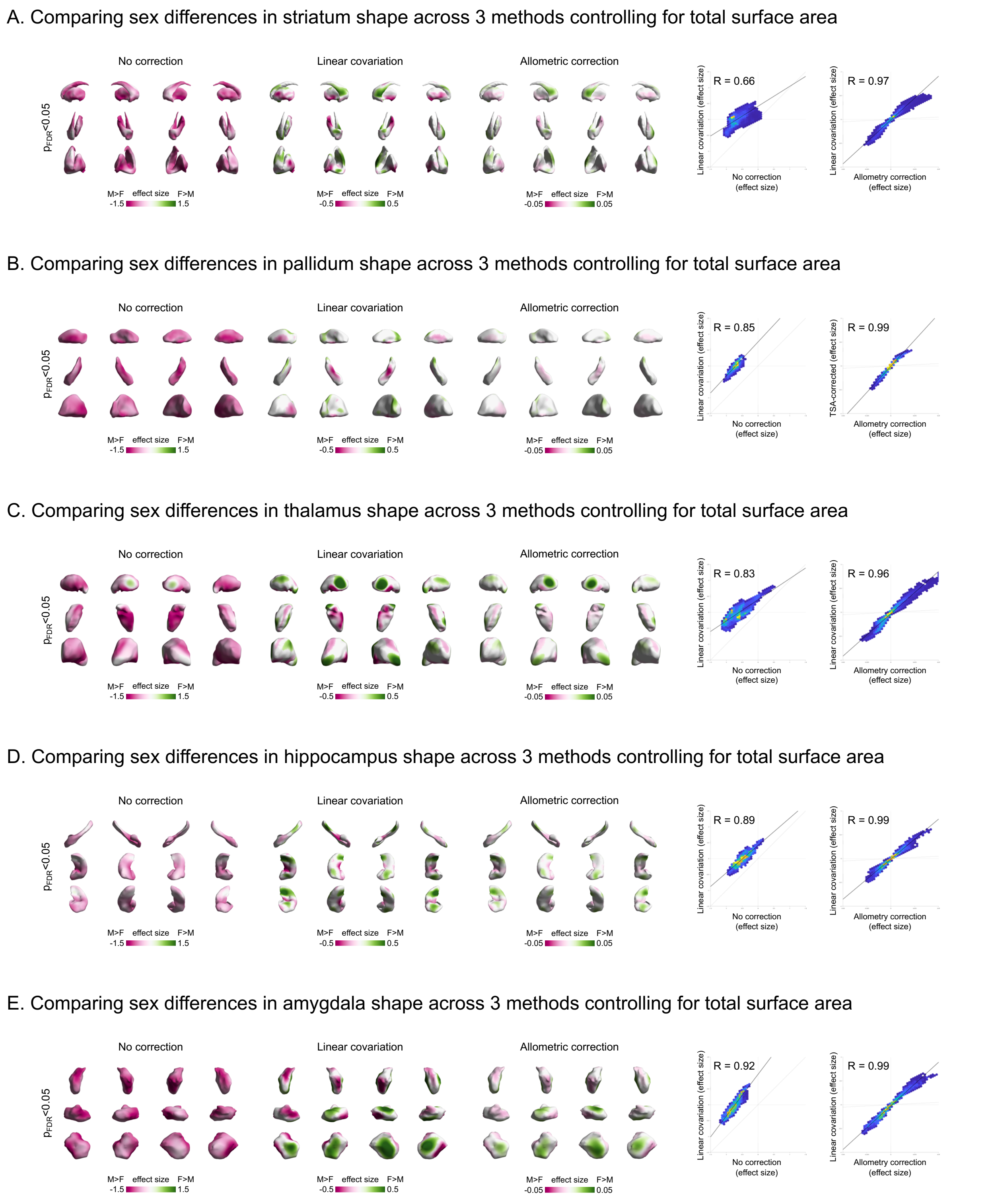


**Supplemental Figure 3. Comparing the sex differences in regional subcortical shape in early adulthood across three approaches to control for total brain size. A.** Standardized effect sizes for sex differences in regional surface area of striatum computed based on no correction (left map), linear covariation (middle map), and allometric correction (right map) of total brain size. Female-biases are shown in green, while male-biases are shown in magenta. The hexplots show structure-wide spatial correction (R) between sex differences computed from no correction vs. linear covariation (left) and between those computed from allometric correction vs. linear covariation. **B.** Same as panel A but for pallidum. **C.** Same as panel A but for thalamus. **D.** Same as panel A but for hippocampus. **E.** Same as panel A but for amygdala.

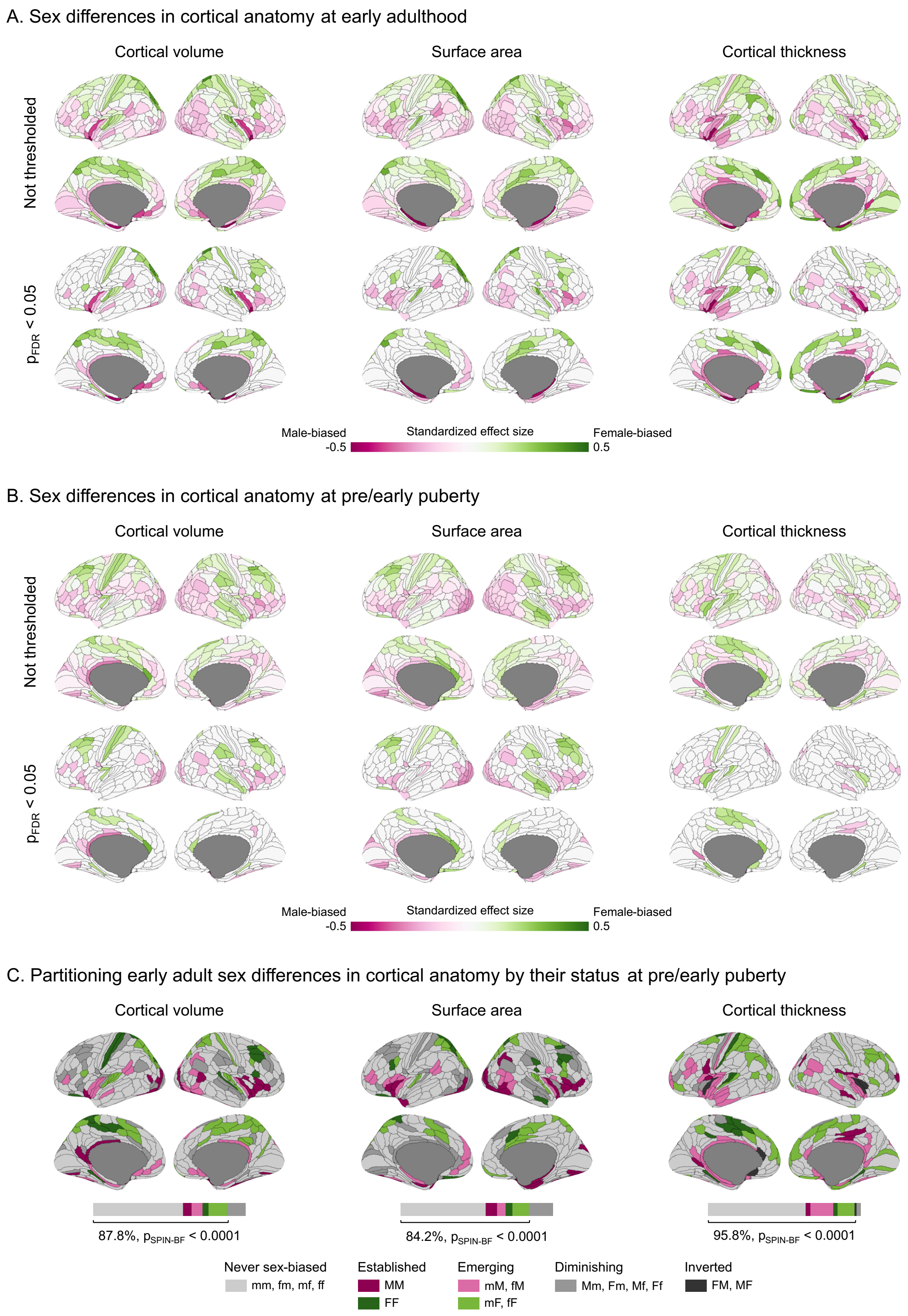

A. Sex differences in cortical anatomy at early adulthood
Cortical volume
Surface area
Cortical thickness
Not thresholded
$p_{FDR} < 0.05$
Male-biased
-0.5
Standardized effect size
Female-biased
0.5
B. Sex differences in cortical anatomy at pre/early puberty
Cortical volume
Surface area
Cortical thickness
Not thresholded
$p_{FDR} < 0.05$
Male-biased
-0.5
Standardized effect size
Female-biased
0.5
C. Partitioning early adult sex differences in cortical anatomy by their status at pre/early puberty
Cortical volume
Surface area
Cortical thickness
87.8%, $p_{SPIN-BF} < 0.0001$
84.2%, $p_{SPIN-BF} < 0.0001$
95.8%, $p_{SPIN-BF} < 0.0001$
Never sex-biased
mm, fm, mf, ff
Established
MM
FF
Emerging
mM, fM
mF, fF
Diminishing
Mm, Fm, Mf, Ff
Inverted
FM, MF

**Supplemental Figure 4. Consistency of sex differences in cortical anatomy between pre/early puberty and early adulthood. A.** Top row shows standardized effect sizes for sex differences (female-biases: green, male-biases: magenta) in regional cortical volume, surface area, and thickness [controlling for age, global phenotypes (total cortical volume, total surface area, or mean cortical thickness), and Euler index] in early adulthood (HCP-YA). The bottom row shows statistically significant sex differences based on a two-tailed t-test followed by false discovery rate (FDR) correction at $p_{FDR} < 0.05$ across 360 cortical regions of the multimodal Glasser parcellation. **B.** Same as panel A but for sex differences in regional cortical anatomy [controlling for age, global phenotypes (total cortical volume, total surface area, or mean cortical thickness), Euler index, and site] in pre/early puberty (ABCD-9Y). **C.** Conjunction maps showing consistency of anatomical sex differences in pre/early puberty (ABCD-9Y) and early adulthood (HCP-YA). Never sex-biased is defined as regions where sex-biases are non-significant in both pre/early puberty and early adulthood (lightest gray). Established is defined as regions where female- and male-biases are significant in the same direction in pre/early puberty and early adulthood (darker green and magenta). Emerging is defined as regions where female- and male-biases are significant in early adulthood but non-significant in pre/early puberty in the same direction (lighter green and magenta). Diminishing is defined as regions where female- and male-biases are significant in pre/early puberty but non-significant in early adulthood (medium gray). Inverted is defined as regions where female- and male-biases are significantly in the opposite directions between pre/early puberty and early adulthood (darkest gray). Bars show the areal proportion of subcortical surface falling within each of these conjunction categories. Brackets and percentages indicate areal proportion of consistent sex differences between childhood and adulthood. $P_{SPIN-BF}$ indicates the statistical significance of consistent areal proportion using a one-tailed spatial permutation test based on 10,000 spin permutations of HCP-YA sex differences (**Methods**). Source data are provided in Source Data - Sex Differences in Human Subcortical Anatomy.xlsx.

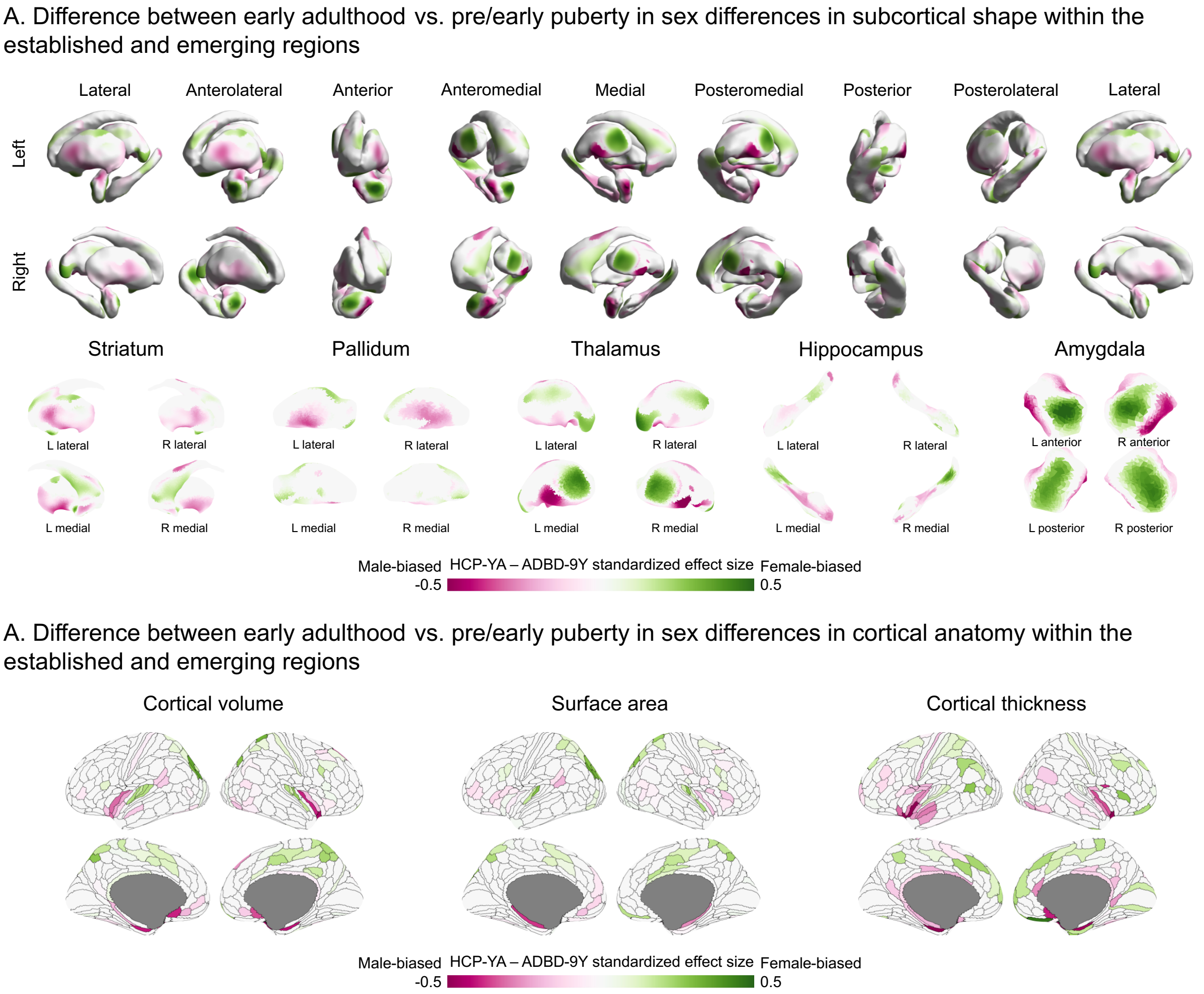


**Supplemental Figure 5. Difference between early adulthood and pre/early pubertal childhood in sex differences in subcortical shape and cortical anatomy within established and emerging regions. A.** The HCP-YA – ABCD-9Y difference in standardized effect sizes for sex differences in regional surface area (controlling for age and total surface area) showing female-biases (green) and male-biases (magenta) within regions found to be established and emerging from Figure 3. **B**. Same as A, but for regional cortical volume, surface area, and thickness [controlling for age, global phenotypes (total cortical volume, total surface area, or mean cortical thickness), and Euler index] from Supplemental Figure 4.

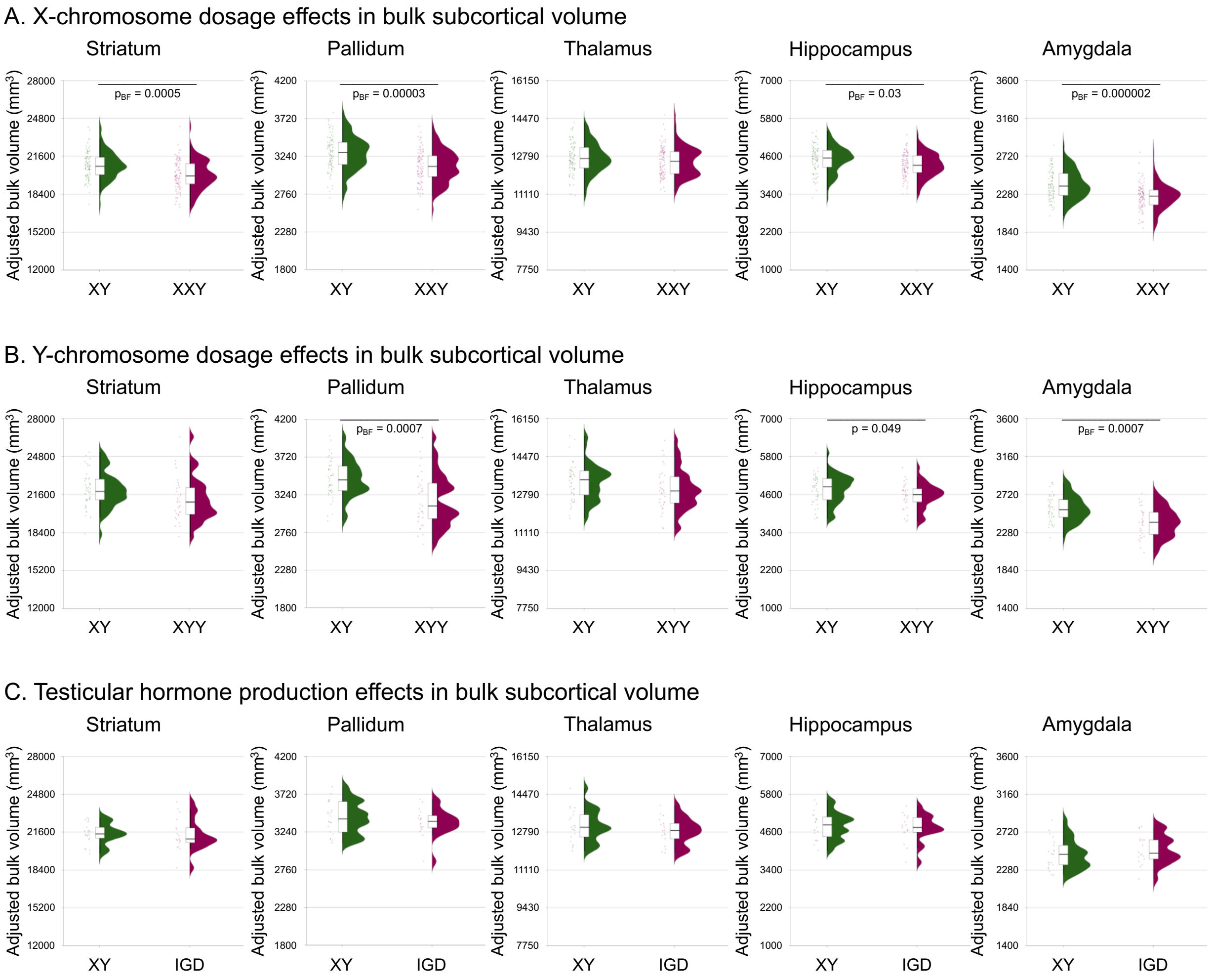


**Supplemental Figure 6. Sex chromosome dosage and testicular hormone production effects on bulk subcortical volume. A.** Bulk volume of subcortical structures (adjusted for age and total brain volume) for XXY (green) and XY (magenta) participants. **B.** Same as panel A but for XYY (green) and XY (magenta) participants. **C.** Same as panel A but for XY males (green) and males with IGD (magenta). For all panels, the boxes, histograms, and jittered dots indicate median bounded by first and third quartiles, distribution, and individual values. Horizontal bars indicate structures showing significant sex differences at nominal $p < 0.05$ or Bonferroni-corrected $p_{BF} < 0.05$.

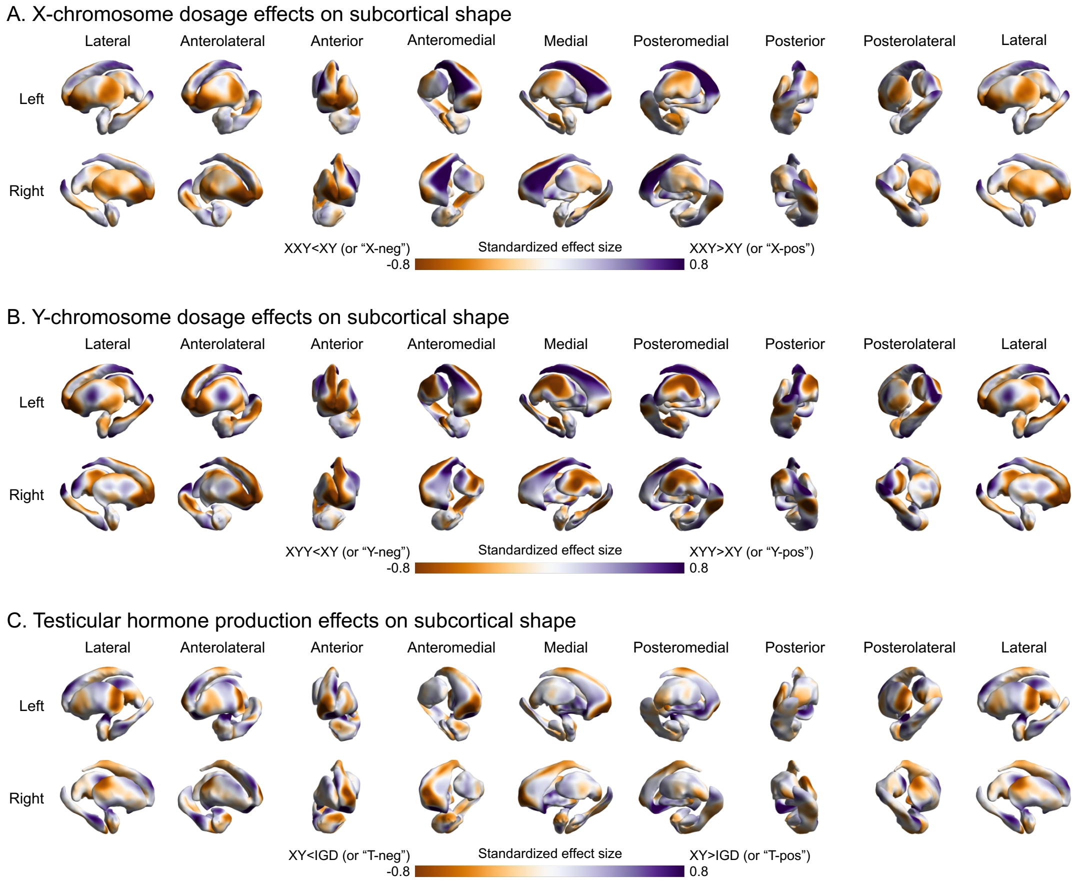


**Supplemental Figure 7. Effects of sex chromosome dosage and testicular hormone production on regional surface area of subcortical structures. A.** Tow row shows standardized effect sizes for X-chromosome dosage effects (XXY vs. XY) on regional surface area (controlling for age and total surface area). Positive effect sizes (XXY>XY or “X-pos”) are shown in purple, while negative effect sizes (XXY<XY or “X-neg”) are shown in orange. **B.** Same as panel A but for Y-chromosome dosage effects (XYY vs. XY). **C.** Same as panel A but for effects of intact testicular hormone production (male controls vs. males with IGD). Source data are provided in Source Data - Sex Differences in Human Subcortical Anatomy.xlsx.

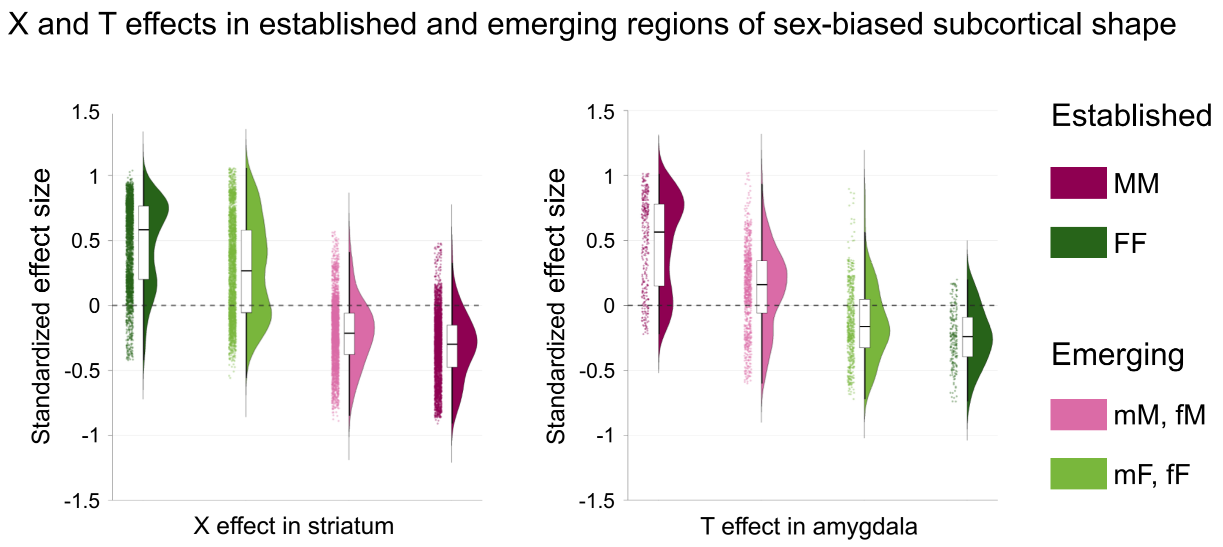


**Supplemental Figure 8. Sex chromosome dosage and testicular hormone production effects within established and emerging regions of sex-biased subcortical shape.** Standardized effect sizes for X-chromosome dosage (XXY vs. XY; "X"), Y-chromosome dosage (XYY vs. XY; "Y"), and testicular hormone production (XY vs. IGD; "T") on regional surface area (controlling for age and total surface area) are plotted for sex-biased subcortical shape. The boxes, histograms, and jittered dots indicate median bounded by first and third quartiles, distribution, and individual values for the X, Y, and T effect sizes. Female-biased regions that are significant in both pre/early puberty and early adulthood (or "established") are shown in darker magenta. Those that are only significant in early adulthood (or "emerging") are shown in lighter magenta. Likewise, established (darker green) and emerging (lighter green) male-biased regions are shown. MM: significantly male-biased in both cohorts; FF: significantly female-biased in both cohorts; mM/fM: significantly male-biased in adult cohort only; mF/fF: significantly female-biased in adult cohort only.
