## Supplemental Tables for "Sex Differences in Human Subcortical Anatomy are Robust and Replicable in Adulthood, Already Evident Prepubertally, and Aligned with Sex Chromosomal and Gonadal Influences"

| | | Sample size (n) | Age (mean ± SD) |
|---|---|---|---|
| **HCP-YA** | **Females** | 592 | 29.56 ± 3.6 |
| | **Males** | 493 | 27.89 ± 3.59 |
| **UKB-MA** | **Females** | 375 | 48.75 ± 0.94 |
| | **Males** | 294 | 48.89 ± 0.83 |
| **ABCD-9Y** | **Females** | 569 | 9.19 ± 0.17 |
| | **Males** | 563 | 9.21 ± 0.17 |
| **SCA XXY** | **XXY** | 99 | 16.38 ± 4.78 |
| | **XY** | 92 | 16.24 ± 5.64 |
| **SCA XYY** | **XYY** | 34 | 15.49 ± 5.33 |
| | **XY** | 47 | 13.97 ± 4.76 |
| **IGD** | **IGD XY** | 19 | 23.27 ± 8.33 |
| | **XY** | 22 | 21.89 ± 7.66 |

**Supplementary Table 1.** Demographic characteristics of Human Connectome Project Young Adults (HCP-YA), UK Biobank Mid Adulthood (UKB-MA), Adolescent Brain Cognitive Development 9-Year-Olds (ABCD-9Y), Sex Chromosome Aneuploidy (SCA), and isolated gonadotropin-releasing hormone (GnRH) deficiency (IGD) datasets.

| | | Striatum (n) | Pallidum (n) | Thalamus (n) | Hippocampus (n) | Amygdala (n) |
|---|---|---|---|---|---|---|
| HCP-YA | **Females** | 591 | 591 | 591 | 591 | 591 |
| | **Males** | 492 | 492 | 492 | 492 | 488 |
| UKB-MA | **Females** | 371 | 371 | 372 | 372 | 372 |
| | **Males** | 285 | 285 | 286 | 286 | 286 |
| ABCD-9Y | **Females** | 568 | 569 | 569 | 568 | 569 |
| | **Males** | 563 | 563 | 562 | 560 | 562 |
| SCA XXY | **XXY** | 91 | 94 | 94 | 91 | 94 |
| | **XY** | 86 | 88 | 87 | 84 | 85 |
| SCA XYY | **XYY** | 31 | 33 | 33 | 32 | 32 |
| | **XY** | 46 | 46 | 46 | 46 | 45 |
| IGD | **IGD XY** | 18 | 18 | 18 | 17 | 18 |
| | **XY** | 22 | 22 | 22 | 20 | 22 |

**Supplementary Table 2.** The number of samples used for analyses after quality control of segmentations for Human Connectome Project Young Adults (HCP-YA), UK Biobank Mid Adulthood (UKB-MA), Adolescent Brain Cognitive Development 9-Year-Olds (ABCD-9Y), Sex Chromosome Aneuploidy (SCA), and isolated gonadotropin-releasing hormone (GnRH) deficiency (IGD) datasets.
